# Rapid Peptide Mapping of Monoclonal Antibodies with Direct Infusion Mass Spectrometry

**DOI:** 10.64898/2026.05.14.725248

**Authors:** Austin Z. Salome, Marcel Morgenstern, Alexander S. Hebert, Craig D. Wenger, Pavel Sinitcyn, Benton J. Anderson, John S. Chlystek, Lia R. Serrano, Keaton L. Mertz, Ian J. Miller, Erik Miller-Galow, Malika P. Godamudunage, Michael Batt, Bhavin R. Patel, Gihoon Lee, Lloyd M. Smith, Scott T. Quarmby, Alayna M. George Thompson, Joomi Ahn, Harsha P. Gunawardena, Joshua J. Coon

## Abstract

Peptide mapping is a critical tool for characterizing biotherapeutic proteins and is essential for the development of monoclonal antibody drugs. Here we describe a new direct infusion technology that streamlines peptide mapping data collection and analysis, accelerating the method by up to 100-fold. This method, which we term RaPiD-mAb-MS, combines high-throughput plate-based sample preparation with direct infusion mass spectrometry analysis. RaPiD-mAb-MS allows analysis of 96 samples within ∼ 1.5 to 2 hours, routinely achieves >95% sequence coverage, and has been successfully applied to 28 unique antibodies and over 2,000 samples. Here we demonstrate that RaPiD-mAb-MS detects and quantifies oxidation, deamidation, isomerization, glycosylation, and sequence variants with results comparable to conventional LC-MS based methods in a fraction of the time. Further, by eliminating chromatography, data analysis is greatly streamlined and simplified. By allowing for the collection of ∼ 1,000 peptide maps per day, RaPiD-mAb-MS is positioned to accelerate all phases of antibody-based drug discovery & development and sets the stage for collection of massive datasets that would allow artificial intelligent prediction of optimal antibody variants and formulations.

## Main

Biotherapeutic proteins have become crucial in the treatment of a wide range of diseases, including cancers, autoimmune disorders, infectious diseases, and genetic and endocrine disorders.^1–8^ These biologic agents, often monoclonal antibodies (mAbs), provide targeted therapies that can exceed the specificity and efficacy of traditional small-molecule drugs. The first mAb approved by the U.S. Food and Drug Administration (FDA) in 1986, Orthoclone OKT3 by Johnson & Johnson,^9, 10^ set the stage for the rapid development of mAb-based treatments and marked the beginning of a new era in biologic medicine. Over the following decades, the use of mAbs has expanded significantly, with the FDA approving its 100th monoclonal antibody 35 years later in 2021.^11^ Since then, the number of mAbs in clinical trials has steadily increased, alongside an overall rise in approval rates.^12^ With 162 FDA approved mAbs and a $200 Billion annual market, mAbs represent a fast-growing and increasingly vital class of medicines.^2^

With the significant increase in the production and development of mAbs, there is a growing need to thoroughly characterize these large molecules at every stage, from initial discovery and development to manufacturing. Throughout these stages, there are numerous opportunities for protein degradation, which can affect both the efficacy and safety of the mAb. Additionally, forced degradation studies are performed to assess overall stability.^13–16^ To address these challenges, a variety of techniques are employed, including liquid chromatography (LC), mass spectrometry (MS), circular dichroism, IR and UV spectroscopy, nuclear magnetic resonance spectroscopy, cryo-electron microscopy, X-ray diffraction, and a variety of immunological and biological assays among others.^17–27^ Among these methods, LC-MS plays a central and pivotal role.^28^

In response to the growing need for more comprehensive and efficient characterization, multi-attribute methods (MAM) have emerged as a powerful analytical approach.^29^ While ’MAM’ is often synonymous with regulated quality control, the core concept is to characterize and monitor multiple product quality attributes (PQAs) simultaneously using a single approach. By allowing for the characterization of multiple PQAs in one method, MAM offers a more holistic view of product quality, enabling more precise control over the manufacturing process and ensuring the final product’s consistency, safety, and efficacy.^30–32^ One such MAM technique is the use of LC-MS for peptide mapping, which is invaluable for both identity assessment and monitoring post-translational modifications (PTMs).^33–37^ Peptide mapping entails LC-MS analysis of enzymatically produced peptides from the target protein. Typically, peptides composing the complete sequence and common chemical and biological modifications (oxidation, deamidation, isomerization, glycosylation, clipping, etc.) are observed. The signals for the modified and non-modified (native) peptides are compared to determine modification occupancies.

These measurements can be correlated with bioactivity/binding analyses to inform strategies for molecular engineering, cell-line development, expression, purification, formulation development, and storage of biotherapeutic proteins to specifically protect critical amino acids.

While LC-MS peptide mapping has extensive utility, there are some major drawbacks to this methodology. Specifically, the collection of peptide mapping data is both cumbersome and complicated, requiring long data collection times and complicated data analyses.^38^ The throughput of the method is inherently limited by the requirement to use chromatography which also increases reagent cost and waste.^39^ Short chromatographic methods can increase throughput, but often these fast separations cannot effectively separate deamidated and isomerized peptides from their native counterparts. Additionally, chromatographic columns must be equilibrated for extended time prior to separation and may require blank runs after every analysis to limit carryover. Although extensively developed, chromatographic peak selection algorithms are not flawless and often require at least a manual check, especially for low level modifications. To mitigate some of these challenges, a variety of automated workflows have been implemented to streamline protein digestion and LC-MS injection; however, the LC-MS data collection time and analysis time remains a bottleneck.^40–46^

Herein we describe a rapid peptide mapping method (Rapid Peptide mapping by Direct infusion - monoclonal antibody-Mass Spectrometry, RaPiD-mAb-MS) for high throughput sample preparation and analysis. RaPiD-mAb-MS eliminates liquid chromatography and instead directly infuses the mAb sample into an high-resolution, accurate mass MS system.^47–50^ Elimination of chromatography affords RaPiD-mAb-MS two major benefits as compared to LC-MS peptide mapping: (1) analysis speeds up to 100 times faster and (2) greatly simplified data analysis. Within approximately 60 seconds, RaPiD-mAb-MS typically achieves >95% sequence coverage and detects modifications including oxidation, deamidation, isomerization, glycosylation, N-terminal cyclization, and the sites of drug conjugation, among others. Here we describe the use of the RaPiD-mAb-MS technology for its ability to characterize a wide variety of unique mAbs (28) and demonstrate it provides results comparable with conventional LC-MS. Finally, we demonstrate the utility of RaPiD-mAb-MS to analyze over 1,000 mAb samples in just 1.5 days, generating a massive quantitative database from which to use machine learning and linear modeling to predict mAb formulations. We conclude that while conventional LC-MS remains key for final quality control due to the entrenchment of FDA-approved methods, RaPiD-mAb-MS breaks the analytical throughput bottleneck essential for accelerating discovery and pre-clinical development.

## Results

### Plate Preparation and Data Acquisition

The first step in development of the RaPiD-mAb-MS direct infusion technology was to create a digestion procedure that eliminates excipients and buffers present in mAb formulations while also preventing loss of short and hydrophobic tryptic peptides. To maximize throughput and compatibility with automation, we utilized a 96 well microtiter plate format. Note, while many peptide mapping sample preparation protocols have been described^40–44, 51, 52^, to our knowledge they are all designed for downstream LC separations and, therefore, require desalting prior to infusion. Our approach accomplishes sample cleanup through an integrated filter plate in which the mAb is reduced, alkylated, precipitated, washed, and trypsin digested (**Figure 1a**). These peptides are displaced from the filter plate to a second standard 96 well microtiter plate by centrifugation in an aqueous solvent. From this plate the peptide can be either directly infused or loaded onto an LC-MS for traditional analysis. In our hands, this protocol is highly effective and eliminates components such as PBS, Proline, Arginine, NaCl, Sucrose, Sorbitol, Trehalose, Citrate, Histidine, Acetate, Tris, Glycerol, PS20, PS80, Tween 20, TCEP, GnHCl, DTT, and IAA.

**Figure 1.**
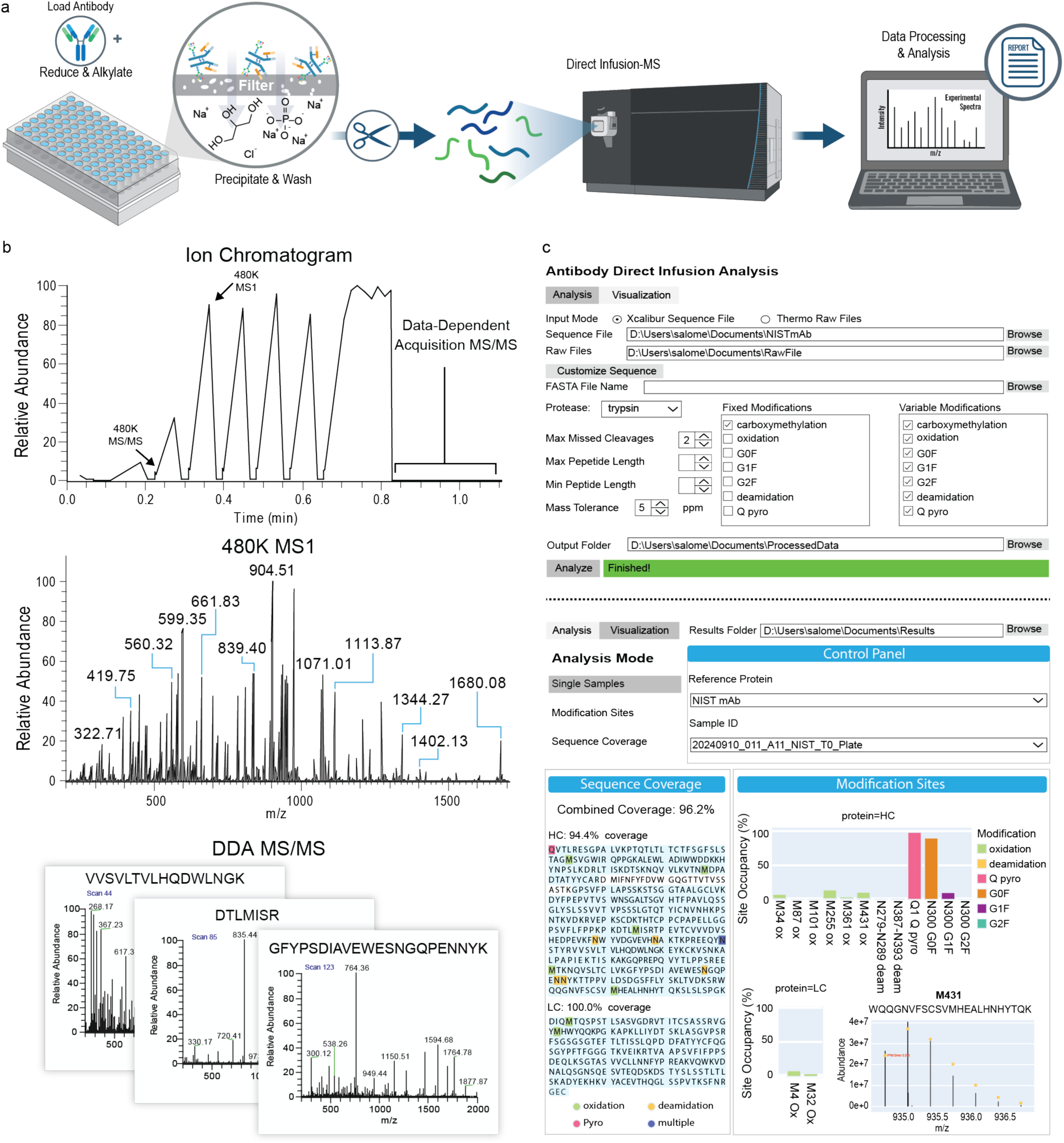
Overview of the RaPiD-mAb-MS methodology. **a**, Overview of sample preparation, direct infusion MS, and data analysis workflow for RaPiD-mAb-MS. Preparation includes reduction & alkylation, precipitation & washing, and digestion followed by direct infusion MS. **b**, Ion chromatogram of a typical RaPiD-mAb-MS method where a mixture of high resolution MS1, targeted MS/MS, and data dependent MS/MS are collected over a timespan ranging from ∼60 seconds. **c**, Overview of custom software that allows for the analysis of RaPiD-mAb-MS data. Once processed, the tool provides visualization capability to examine sequence coverage, modifications, and raw MS data.

These samples are then ready for direct infusion into an MS system. We found that nano-electrospray (nESI), as compared to the more conventional micro-electrospray (µESI), minimized the amount of in-source oxidation. nESI is easily automated by use of the Advion NanoMate system wherein each sample is sprayed with an individual, single use ESI nozzle. The NanoMate was then coupled with a hybrid Orbitrap MS system (i.e., Eclipse or Astral) that have the ability to achieve high resolution accurate mass measurements. We utilized a scan sequence (**Figure 1b**) that mixed high resolution MS1 (480K @ *m/z* 200), high resolution MS/MS (480K, for targeted deamidation analysis), and lower resolution data-dependent MS/MS (∼15-80K). In its entirety, this MS method typically takes ∼60 seconds per sample, so that sample-to-sample cycle times are approximately 120 seconds (i.e., method time plus sample changeover).

**#Supplementary Figure 1** presents an MS1 scan from the direct infusion of NIST mAb prepared as described above. By calculating the theoretical *m/z* values of the *in silico*-generated NIST mAb tryptic peptides, we identify a corresponding *m/z* peak to within 10 ppm for all but one of the predicted tryptic peptides. The single unobserved species (DMIFNFYFDVWGQGTIVTVSSASTK, residues 375-399) is a highly hydrophobic heavy chain CDR3 peptide. We attribute its absence to solubility limits in the aqueous spray solvent, as subsequent tests adding organic solvent to the infusion solution enabled its detection. Despite this specific outlier, the result is a sequence coverage of 96%, where even very short peptides that would typically not be retained during LC-MS are observed (e.g. VSNK)^51, 53^. To automate this process, we developed software (RaPiD-Analyze) which performs an *in silico* digest using input antibody protein sequences and generates matches based on user defined mass and isotopic pattern error tolerances (**Figure 1c**). In our experience, the simplicity of these mixtures allows for high confidence identification based on these features alone; however, MS/MS scans can also be used to increase ID confidence. RaPiD-Analyze also calculates the site occupancy of modifications by taking the ratio of the isotopic cluster abundances for the modified form divided by the sum of all forms of the same peptide (including various charge states).

To determine the optimal data collection settings for these direct infusion experiments we examined several parameters including sample concentration, number of ions to inject (i.e., AGC targets), and resolving power (**Supplementary Figure 2**). We selected the following baseline parameters: 4.5 µM sample concentration, 1-2.5 million AGC target, and 480,000 resolving power. Note we see strong performance at all resolving powers above ∼ 100,000. The MS1 scan provides valuable insights into modifications like oxidation and glycosylation amongst others.

Having developed a sample preparation protocol, instrument methodology, and data analysis software we next sought to evaluate overall method performance. First, we tested sensitivity with eight stable-isotope labeled NIST mAb peptides that were spiked into a NIST mAb digest at varying concentrations. Using these data we calculated the limit of detection (based on 3X spectral signal-to-noise) for each and used the values to determine the lowest ratio at which the labeled peptides could be detected – simulating the ability to detect low level modifications and sequence variants, for example. On average, the method detects peptides as low as ∼ 0.7% relative abundance, with values ranging from 0.2% - 1.2% (**Extended Data Figure 1**). The variation likely depends primarily on the ionization efficiency of the individual peptide sequence. Note preliminary work with selected ion monitoring (SIM) or other targeted strategies hold promise to further reduce detection limits.^54^

Obtaining high sequence coverage is a key Figure of Merit for peptide mapping experiments. Owing to its good sensitivity, we supposed that the RaPiD-mAb-MS approach would allow for high sequence coverage. To test this hypothesis, we obtained 24 antibodies derived from three main base sequences of Clesrovimab, Nirsevimab, and Semorinemab. As expected, we achieved very high sequence coverage – 100% of the sequence for all Nirsevimab and Semorinemab sequences (118 and 125 unique peptides, respectively) and ∼ 95% coverage of the Clesrovimab molecules (127 unique peptides) (**Extended Data Figure 1k**).

### Detection of modifications using MS1

Unlike conventional peptide mapping where the site occupancy of any given modification is calculated by determining the area under the chromatographic peak (for each species), the RaPiD-mAb-MS approach simplifies this process by directly using *m/z* peak intensities. **Figure 2a** illustrates this process with the notoriously oxidized DTLMISR sequence. From the MS1 scan we observe *m/z* peaks at 835.43 and 851.43, corresponding to the singly protonated native and oxidized versions of this peptide. The inset displays the isotopic clusters and their relative intensities which are then used to calculate a relative percent modification occupancy – 19.6%.^30^ Next, we sought to compare quantitative performance of RaPiD-mAb-MS to conventional LC-MS peptide mapping by subjecting NIST mAb to forced degradation. The results of these experiments are shown in **Figure 2b**, where excellent correlation is observed for numerous peptides with methionine and tryptophan oxidation (r^2^ > 0.99). To ensure similar performance across antibodies with varying sequences we performed the same experiment with Sigma mAb, Infliximab, Rituximab, and Cetuximab with single timepoints and the complete dataset shown in **Extended Data Figure 2**. Again, we note excellent correlation (r^2^ > 0.99) with only slight disagreement at extremely high levels of occupancy between RaPid-mAb-MS and LC-MS. Interestingly, the RaPiD-mAb-MS approach has better precision.

**Figure 2:**
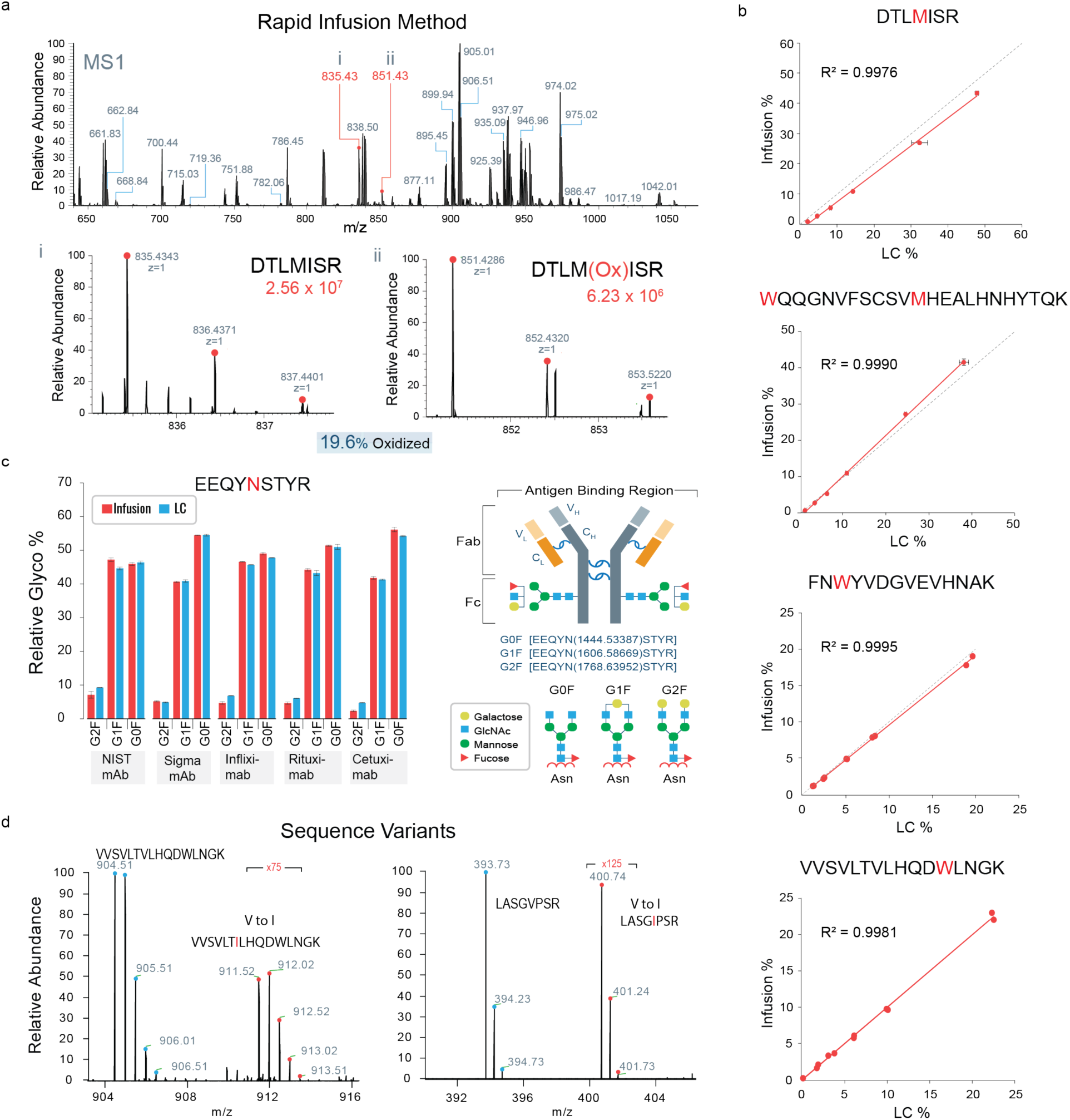
Comparison of RaPiD-mAb-MS to LC-MS peptide mapping. **a,** RaPiD-mAb-MS MS1 spectrum showing the m/z peaks that form the isotope cluster of the (i) native and (ii) oxidized DTLMISR peptide and how these m/z peaks are used to calculate abundance and relative modification occupancies. **b**, Comparison of RaPiD-mAb-MS and LC-MS quantification measurements for oxidation of M and W containing NIST mAb peptides (n=3, dashed line is y=x). Linearity (R^2^) in the 0-10% occupancy range for the peptides shown is summarized in **Supplementary Table 5**. Note that several similar plots of modifications in other mAbs can be found in **Extended Data** Figure 2. **c,** Glycosylation % determination using either RaPiD-mAb-MS and LC-MS (n=3) for the well-known G0F, G1F, and G2F glycoforms of the EEQYNSTYR peptide across five mAbs. Additionally, the structure of the different glycoforms is shown. **d**, RaPiD-mAb-MS measurements of two known sequence variants in NIST mAb samples. Here, each of these sequence variants is readily detected in the MS1 scan at ∼0.7% occupancy.

Glycosylation is another common modification on mAbs and these critical modifications are likewise observed in RaPiD-mAb-MS MS1 data. We evaluated the relative abundance of the well-known G0F, G1F, and G2F on the five mAbs described earlier with both RaPiD-mAb-MS and LC-MS, again finding excellent agreement (**Figure 2c**). Finally sequence variants due to amino acid misincorporations in mAb generation are common and profiled using peptide mapping. We manually searched for the presence of known sequence variants in the NIST mAb data and identified two V to I variants, one at 0.67% and the other at 0.71% respectively. Although both I and L are possible products, the most likely product due to G/U mismatch or misacylation is reported^55^ (**Figure 2d**). Similarly, the method detects and quantifies C-terminal lysine clipping. In NIST mAb, we consistently observe high clipping on the heavy-chain terminal peptide SLSLSPGK (∼90% occupancy), in excellent agreement with LC-MS measurements.

### Detection of modifications using MS/MS

Deamidation is another common modification that occurs in proteins wherein the amino acid asparagine (N) is converted to aspartic acid (D), increasing the mass by 0.984 Da. The subtle change in mass, and chemical structure, adds complexity to the conventional LC-MS peptide mapping method, requiring longer gradients and separation of these distinct peptides. **Figure 3a** shows the chromatographic separation of GFYPSDIAVEWESNGQPENNYK (referred to as PENNYK henceforth) from its deamidated versions. The lower panel of **Figure 3a** presents the isotopic clusters for the native and deamidated forms of PENNYK. Note the isotopic overlap, wherein the first ^13^C isotope of the native peptide and the monoisotopic peak of the deamidated species are separated by only 19 mDa. For RaPiD-mAb-MS, where no chromatography is performed, native and deamidated peptides have isotopic *m/z* clusters that overlap and are not always separable at 480K MS1 resolving power. In such cases, we perform a targeted 480K resolving power MS/MS scan on the native/deamidated peptide precursor population to generate product ions having low enough *m/z* and charge such that the isotopic clusters are resolved, and quantification can be achieved (**Figure 3b**). Note that since any given antibody contains only ∼12-17 Asn-containing peptides, we generate a comprehensive list of all possible deamidated peptides for *a priori* targeting. In this way, the method functions as a discovery tool, since interrogating this complete list adds only ∼15 seconds of overhead to the analysis.

**Figure 3.**
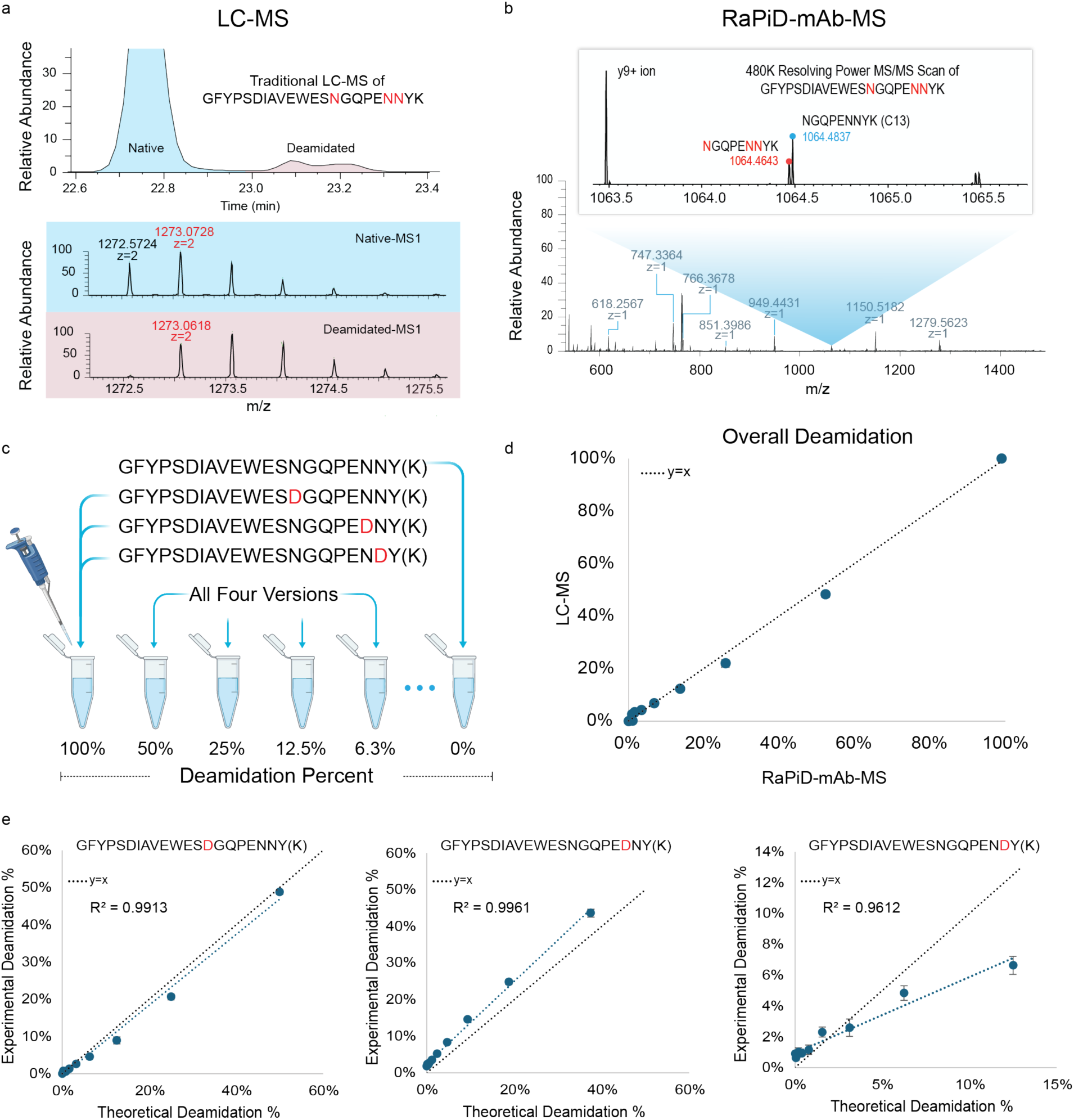
Approaches for quantifying deamidation by conventional LC-MS and RaPiD-mAb-MS. **a,** Because they are so similar in mass, conventional LC-MS peptide mapping relies on chromatographic separation of deamidated peptides from their unmodified counterparts. The chromatographic areas of each separated species is then summed and used to determine overall modification occupancy. **b**, RaPiD-mAb-MS utilizes the high resolving power of Orbitrap MS systems to directly distinguish the deamidated from unmodified peptide. This measurement is made during an MS/MS scan so that the site of deamidation can be localized to an individual residue. **c**, Titration of heavy ^13^C labeled native and deamidated PENNY(K) peptides to make solution from 100% deamidated to 0% deamidated in a background of NIST mAb peptides. **d**, Comparisons of LC-MS and RaPiD-mAb-MS analysis of the spike-in mixtures showing high correlation between the methods (n=3). **e**, Site specific titration of each deamidation site on ^13^C labeled PENNY(K) peptide. Linearity (R^2^) in the 0-10% occupancy range for the peptides shown in panels d and e is summarized in **Supplementary Table 5**. Note that related data on FNWYVDGVEVHNA(K) can be found in **Extended Data** Figure 3.

To test the performance of this strategy we synthesized stable isotope labeled versions of PENNYK and FNWYVDGVEVHNAK with each N fully deamidated. These synthetic peptides were mixed at known ratios with an additional non-deamidated labeled version as shown in **Figure 3c**. These mixtures were then spiked into the NIST mAb background and analyzed using both RaPiD-mAb-MS and LC-MS. **Figure 3d** presents the comparison for overall deamidation measurement of these methods for PENNYK (see **Extended Data Figure 3** for FNWYVDGVEVHNAK). From these results we conclude the targeted approach is comparable in performance to LC-MS for overall deamidation quantification. Next, we utilized the product ion isotopic abundances to localize the deamidation modification to a specific residue within the peptide sequence, seeing how well the experimental ratios matched. **Figure 3e** demonstrates that we can localize quite confidently for the two Ns on the N-terminal side; while the C-terminal N measurement is linear, it has less accuracy. This is not entirely surprising since distinguishing between the two sequential Asp residues affords fewer product ions with which to obtain measurements. Overall, however, we conclude that the use of high-resolution MS/MS scanning can allow for accurate deamidation abundance estimates and reasonably good site localization (we believe further algorithm development will likely improve this capability).

### Detection of antibody-drug conjugates

Antibody-drug conjugates (ADCs) are an increasingly important class of biotherapeutics that are also characterized using peptide mapping. To test the RaPiD-mAb-MS approach for ADCs, we obtained two humanized Immunoglobulin G (IgG) ADC molecules from Genentech (Genentech IgG 1 and 2). Genentech IgG 1 has a non-cleavable MPEO cystine linker that attaches the drug, a derivative of maytansine (DM1) (**Figure 4a**). Analysis of this molecule using RaPiD-mAb-MS produced the MS1 spectrum shown in **Figure 4b** where it is easily observed that the peptide having the sequence KTCPSSLGQHTVECAYVKH appears at an *m/z* value of 1079.4773, corresponding to the triply protonated version that also contains the MPEO+DM1 modification. During the data-dependent acquisition portion of the RaPiD-mAb-MS method the MS/MS scan of the precursor at m/z 1079.4773 produces the spectrum shown in **Figure 4c**. From these data we confirm the sequence above, observe the product ion for DM1 (*m/z* 547.22), and localize the drug and linker to C-terminal cysteine. **Figure 4d** shows the DM1 immonium ion (*m/z* 547.22) detected across all collected DDA MS/MS scans. Six of these scans contain the signature product ion for DM1, each also mapping to a peptide (e.g., missed cleavage, alternate charge state, and water addition) that contains the previously shown cysteine. Given the modification is only detected at a single site, our data aligns with the known average drug-antibody ratio (DAR) value of 2.^56^ **Figure 4e-h** presents the same analysis but performed on Genentech IgG 2 – a molecule that attaches DM1 to lysine residues with a non-cleavable maleimidomethyl cyclohexane-1-carboxylate (MCC) thioether linker. We note here similar results to the previous example, however, now we localize the DM1 molecule to six different lysine residues. Given this modification is occurring at numerous lysine residues, these data again align well with the provided DAR value of 3.6.^56, 57^ We note these data confirm known chemistries of these molecules as Genentech IgG 1 had site specific Cysteine mutations to allow for reaction of linker and drug with much higher specificity whilst the lysine attachments tend to be more promiscuous.

**Figure 4.**
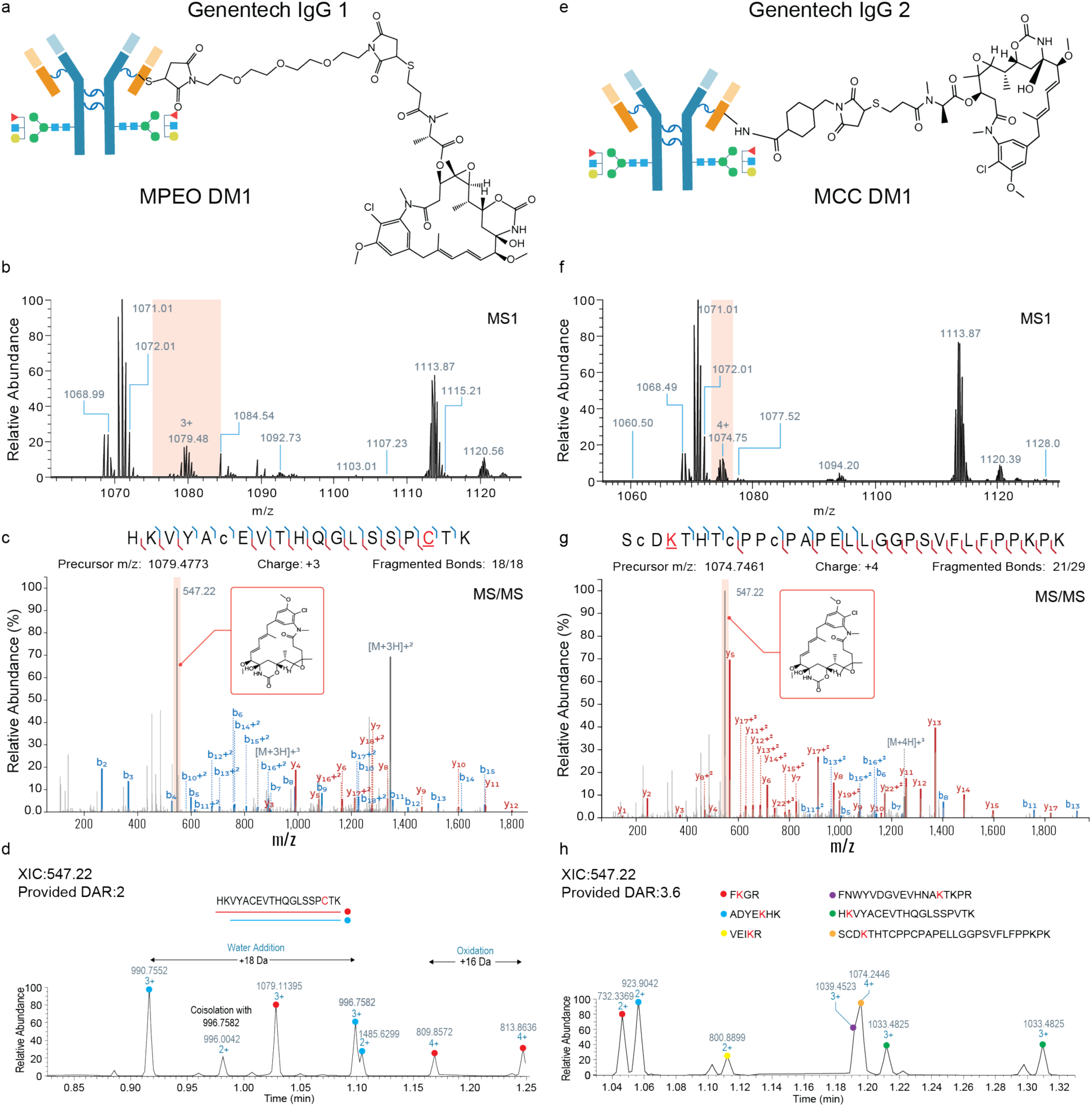
Use of RaPiD-mAb-MS for analysis of Antibody-Drug Conjugates (ADCs). **a**, Linker and drug structure for Genentech IgG 1. **b,** MS1 region of RaPiD-mAb-MS infusion data highlighting the isotope distribution of an intact peptide with MPEO linker and DM1 drug present, as confirmed by accurate mass. **c**, MS/MS spectrum of the identified MS1 peak at m/z value 1079.48 (HKVYACEVTHQGLSSPCTK + MPEO and DM1). This MS/MS spectrum both confirms the accurate MS1 identification and allows for localization of the drug and linker to the underlined cysteine. **d**, Extracted immonium ion (*m/z* 547.22) detected across all DDA scans. Here we identify numerous MS/MS scans that contain evidence of the drug, software analysis confirms that each of these result from other charge states of the above peptide and one missed cleavage peptide. **e-h**, Shows the same analysis but for Genentech IgG 2, a MCC DM1 peptide linker. Drug-antibody ratio (DAR) values were provided by Genentech.

To further test the RaPiD-mAb-MS approach for ADCs we obtained an additional IgG ADC from AbbVie (AbbVie-AB095-PZ). This molecule, which has previously been analyzed, has a surrogate drug and an MC-VC-PAB linker that binds with cysteine (**Extended Data Figure 4a**). Use of RaPiD-mAb-MS for this molecule generates a similar MS1 spectrum where the peptides containing the linker and drug are easily detected based on *m/z* values (**Extended Data Figure 4b**). **Extended Data Figure 4c** presents an MS/MS scan of the precursor at m/z 1,216.6245 and having the sequence THTCPPCPAPELLGGPSVFLFPPKPK. From these data we localize the drug surrogate to the C-terminal side cysteine. A benefit of eliminating chromatography is that short, hydrophilic peptides that normally would not be retained during LC-MS are readily observed. In this analysis, we provide evidence that the drug is also attached to cysteines in the tryptic peptides having the sequence GEC and SCDK (**Extended Data Figure 5**). Previous publications describing the LC-MS characterization of this molecule describe the challenges of conventional methods to observe these known sites.^58^

### Comparison of RaPiD-mAb-MS with conventional LC-MS peptide mapping

All the previous comparisons of RaPiD-mAb-MS to LC-MS peptide mapping were performed within our university laboratory (by AZS and ASH). To directly compare the new technology to industry state-of-the-art, we utilized RaPiD-mAb-MS to analyze a set of 275 forced degradation NIST mAb samples that had been prepared and previously measured at AbbVie (by A.M.G.T. and M.P.G.). RaPiD-mAb-MS samples were prepared using the methods as described above and were collected within approximately 12 hours. The Abbvie LC-MS results were obtained by using a standard protocol that mainly differs from the RaPiD-mAb-MS protocol by its incorporation of a resin-based desalting step (see methods for details). Resultant peptides were separated using a 40 minute, high-flow LC-MS method on an Orbitrap Q-Exactive Plus MS system, followed by data analysis using Byos protein characterization software (eight days LC-MS analysis with one day data searching).

We first calculated how well correlated each of the modification occupancy estimates were between RaPiD-mAb-MS and LC-MS methods. **Figure 5a** shows the difference (in percent) between the two measurements for methionine oxidation of four unique peptides across all 275 samples. Here we see strong correlations with mean differences approaching zero for each. Detailed results for a subset of the data are shown in panels b and c of **Figure 5**. Here we observe that each method detects identical trends and that the results are generally within a percent difference. We note RaPiD-mAb-MS generally measures slightly lower oxidation levels, but expect this subtle difference is most likely an artifact of sample preparation. To provide a global view of these oxidation measurements we constructed heatmaps where each cell is the measured oxidation occupancy (percentage) for the DTLMISR peptide (**Extended Data Figure 6a**). From these results we conclude the results are in excellent agreement across these forced degradation conditions.

**Figure 5.**
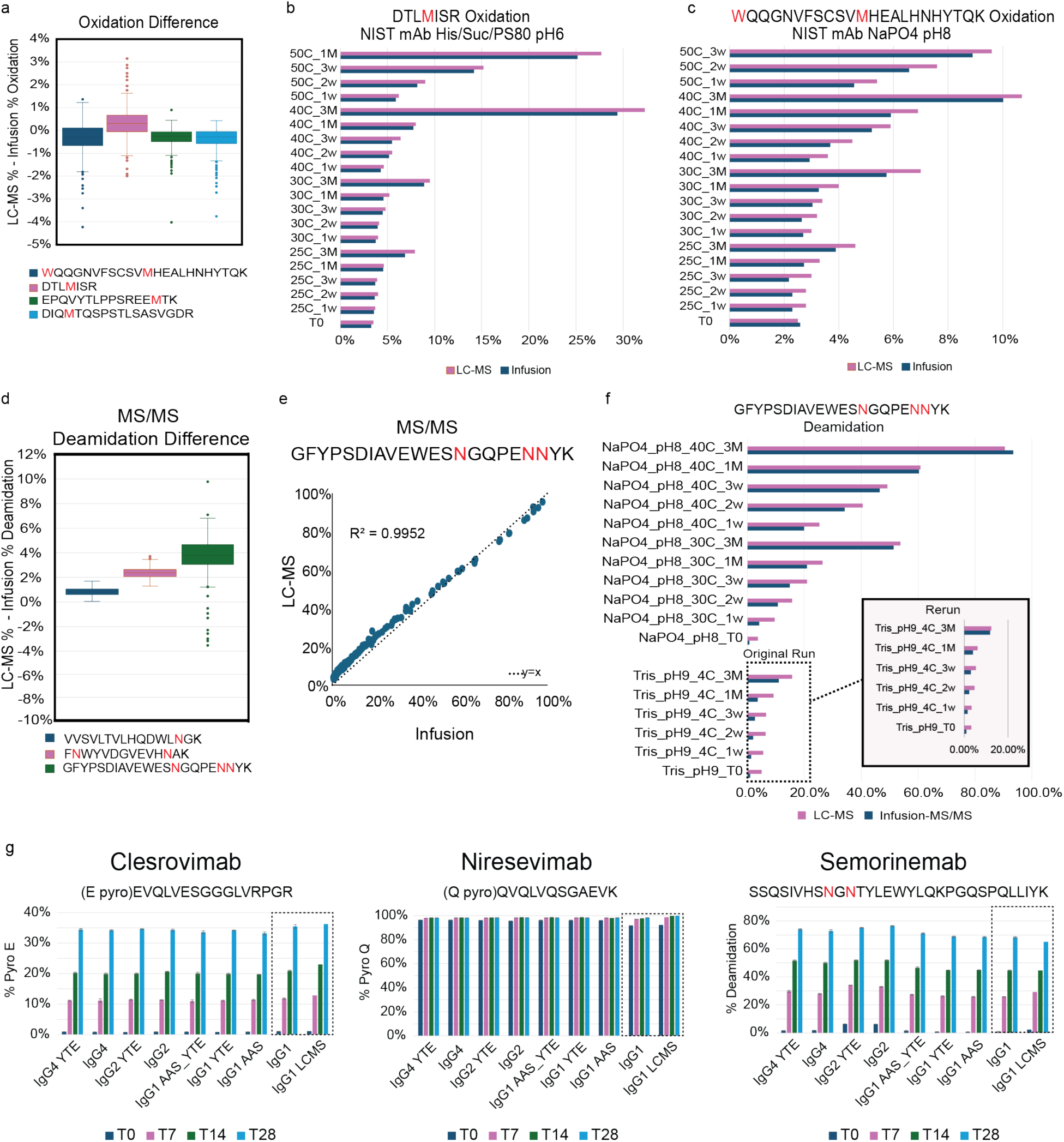
RaPiD-mAb-MS and LC-MS peptide mapping comparison. **a**, Difference (in percent) between the two methods for relative quantification of methionine oxidation of four unique peptides across all 275 samples. A subset of detailed data for selected peptides (**b-c**). **d**, Difference (in percent) between the two methods for relative quantification of asparagine deamidation. A subset of detailed data for the PENNYK peptide (**e-f**). Inset (f) shows the reanalysis using LC-MS of the RaPiD-mAb-MS prepped samples, confirming that the initial difference was largely being driven by deamidation that was occurring in the conventional LC-MS sample preparation. **g**, RaPiD-mAb-MS and LC-MS comparison of a variety of mAbs across time points for N-terminal cyclization and deamidation. **Extended Data** Figure 7 provides additional comparisons for this dataset. All values shown represent peptide-level (summed) occupancies rather than residue-localized measurements.

**Figure 5d** shows the difference between the two measurements for asparagine deamidation across three different modification sites of all 275 samples. Here we see the RaPiD-mAb-MS method is measuring a slightly lower level of deamidation (∼1-3%) as compared to the LC-MS result. This was surprising so we plotted the values each method recorded for the PENNYK peptide (panels e and f of **Figure 5**). Again, we have excellent correlation with the lower level deamidation samples showing the highest disagreement. To determine the source of the discrepancy a subset of RaPiD-mAb-MS prepared samples were reanalyzed using the LC-MS method. These experiments generated an overall lower deamidation occupancy calculation (**Figure 5f**, inset), confirming deamidation levels were likely increased during the AbbVie digestion protocol. **Extended Data Figure 6b** provides the original PENNYK deamidation occupancy data across all forced degradation conditions for both methodologies – again we see identical trends between the two measurements. We note that the RaPiD-mAb-MS technology required only 12 hours to collect and process while the LC-MS approach was nine days.

To evaluate performance across a variety of mAbs and modification sites we obtained 24 unique mAbs (Eli Lilly), each derivative of three primary molecules (clesrovimab, niresevimab, and semorinemab). These mAbs were subjected to forced degradation and then analyzed using both RaPiD-mAb-MS and conventional LC-MS. **Figure 5g** highlights exemplary modifications – N-terminal cyclization (E/Q pyro) and very high level deamidation. We observe time-dependent increases in these modifications and demonstrate the ability to monitor both low- and high-occupancy modifications. Note for IgG1 we performed LC-MS analysis to provide a comparison (see boxed region in each plot). We examined these trends across all molecules and other peptides and observed similar results (**Extended Data Figure 7**).

### High-throughput forced degradation analysis of 1000+ mAb samples

As a final evaluation and large-scale test of the RaPiD-mAb-MS technology, we performed a forced-degradation experiment at Johnson & Johnson using NIST mAb, belimumab (Benlysta), nivolumab (Opdivo), and an additional humanized IgG (MAB1). **Figure 6a** and **Supplementary Tables 1-3** provide an overview of the experimental design that includes three plate conditions (control, and two stressors: high temp and photo), for the four mAbs, and two distinct experiments (buffer and oxidation) for 96 unique combinations per plate. Each of the four mAbs thus has 288 unique conditions spanning three plates and resulting in a total of 12 plates (1152 samples). Briefly, the buffer experiments utilized all two-way combinations of four buffers (citrate @ pH 3.5 & 4.5, acetate @ pH 4.5 & 5.5, histidine @ pH 5.5 & 6.5, and tris @ pH 7.0 & 9.0) and five excipients (arginine, sodium chlorine, proline, sorbitol, and sucrose). The oxidation experiments used four buffers (citrate @ pH 4.0, acetate @ pH 5.2, histidine @ pH 6.0, and tris @ pH 8.0) and many combinations of peroxide, methionine, iron, and EDTA. For each mAb, one plate was used as control, one was heated to 40°C for two weeks, and one was exposed to photons for 30 Whrs.

**Figure 6.**
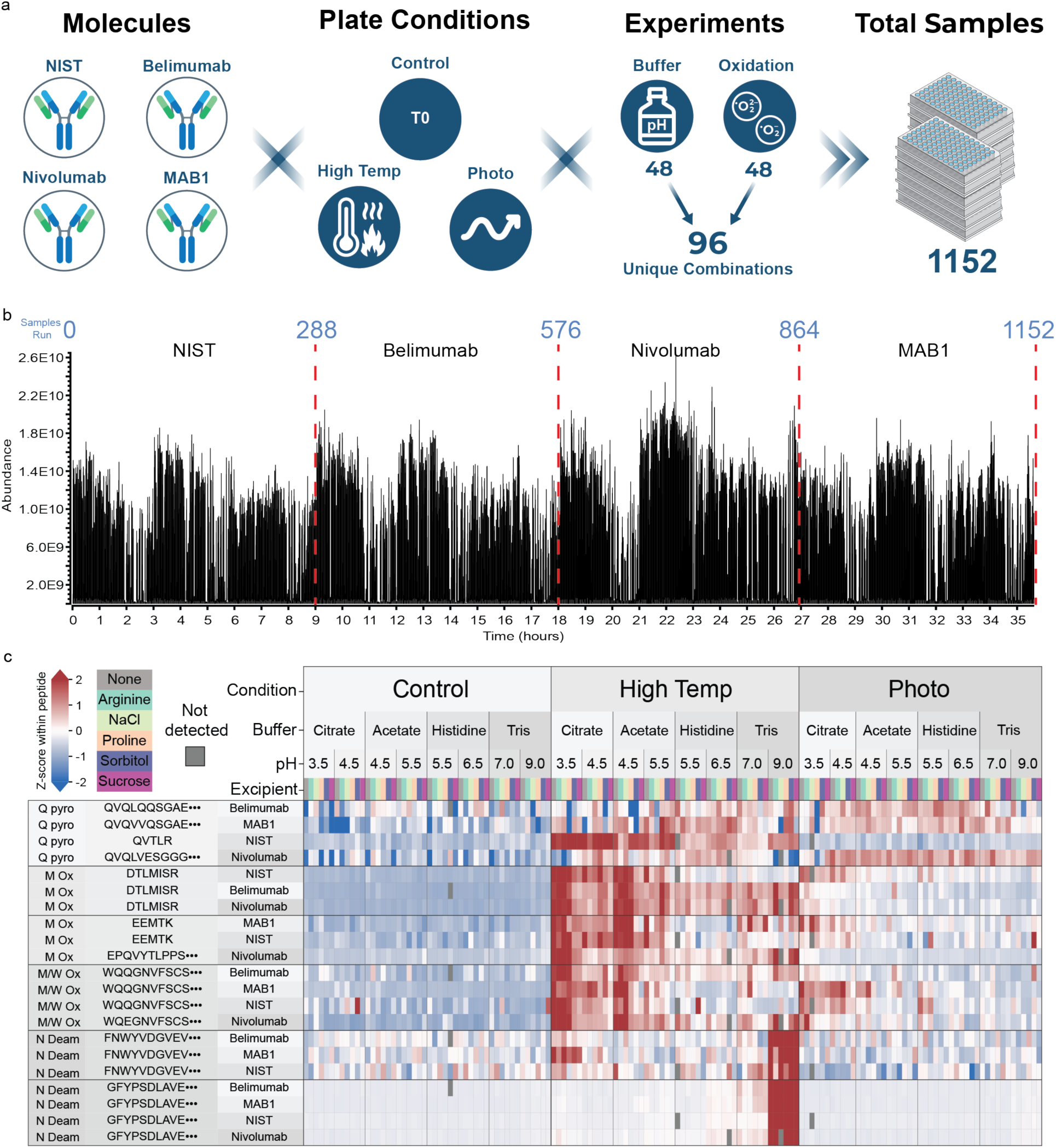
Experimental design and RaPiD-mAb-MS results for analysis of 1152 sample cohort. **a**, Outline of the plate conditions (Control, High Temp, and Photo), mAbs, experiment conditions (oxidation vs buffer), and overall samples analyzed using RaPiD-mAb-MS. Specific condition information can be found in **Supplementary Tables 1-3**. **b**, Combined total ion current plot of the RaPiD-mAb-MS experiments. Each spike represents an individual sample infusion, and the x-axis shows the elapsed time (∼35.5 h). **c,** Heatmap showing the measured quantitative differences for the conserved modifications sites across four antibodies for the 144 buffer experiment conditions.

These twelve plates were prepared and analyzed using the RaPiD-mAb-MS method and the resulting total ion chromatogram is presented in **Figure 6b**. Of the 1152 samples, 1077 resulted in sequence coverage > 96%, with belimumab, nivolumab, MAB1 all yielding ∼ 100% coverage. 75 of the 1152 samples, however, produced lower than expected sequence coverage. Of these, nine had no or very little starting sample and the remaining 66 resulted from the high iron concentration samples and had visible iron chunks following digestion. We conclude that when very high amounts of iron are used a buffer exchange should be performed prior to sample preparation. We note the analysis of these samples was achieved in 35.5 hours while data analysis required an additional four hours.

**Figure 6c** displays a birds-eye view of the buffer experiment displaying modifications observed on the conserved peptides across the four mAbs. In this heatmap the conditions are ordered on the columns while the conserved peptides are grouped on the rows. Because the modification occupancy varies greatly from zero to nearly 100%, displaying the raw percentage would fail to capture the subtle but significant low-level changes as they would be masked by the most abundant ones. Therefore, we applied a z-score where the values for each peptide are converted to standard deviations and then assigned a color to indicate whether each condition is above (red) or below (blue) the average. To illustrate the benefit of this approach, consider the two deamidation sites – one on PENNYK and the other FNWYVDGVEVHNAK – each have increased levels of deamidation in tris pH 9.0 buffer at high temperatures; however, PENNYK is ∼ 40% deamidated whilst FNWYVDGVEVHNAK is ∼ 1%. As expected, with high temperature (40°C) we observe an overall increase in modification occupancy, both within and across mAbs. For example, with citrate and acetate buffers, we observe a trend towards higher oxidation at low versus high pH of methionine.

**Extended Data Figure 8** presents a second heatmap containing the results of the oxidation experiment, again for conserved peptides across all four mAbs. Modified peptides are presented along the rows while across the columns are the various conditions including addition of EDTA, iron, peroxide, methionine, and various combinations thereof. High temperature and high pH cause increases in deamidation, reproducing our observations from the buffer experiment. A notable trend is that methionine oxidation is easily increased by the addition of iron and peroxide; however, the addition of free methionine does an excellent job limiting this modification in the presence of peroxides. Peroxides and high temperatures increase methionine oxidation considerably more than tryptophan oxidation. Tryptophan and methionine are readily oxidized by photons and iron, while the presence of EDTA curbs the modification. **Extended Data Figure 9** presents a subset of these data looking at both M and W oxidation on a subset of NIST mAb peptides having M (EEMTK), W (VVSVLTVLHQDWLNGK), and M and W (WQQGNVFSCSVMHEALHNHYTQK). Here, in a more familiar format, we observe how peroxides affect M but not W, photons and the presence of certain buffers mainly drives W oxidation, and peptides having both M and W are sensitive to both stressors. Further iron appears to drive oxidation on both residues, but when combined with photons, the process escalates. A more comprehensive list of NIST mAb modifications and their abundances across the various conditions is presented in **Extended Data Figure 10**.

### Large-scale data with machine learning can guide mAb formulation

Given the ability of the RaPiD-mAb-MS technology to accelerate the generation of quantitative peptide mapping data, we supposed this method could also provide the sufficiently large datasets required to train artificial intelligence (AI) models. We tested this hypothesis using the data generated with the aim of identifying conditions that minimize modifications on NIST mAb. Starting from a linear machine learning model, we could quickly determine the ideal conditions that would minimize all quantified modifications. The model determined the ideal conditions were Histidine buffer at pH 6.5 with Proline excipient (**Extended Data Figure 10c)**. The model also informed important factors about the effects of the various conditions on performance, for example, using histidine buffer at pH 6.5 led to more than 1.6 percentage point decrease in overall modification rate compared to the worst buffer-pH combo of Tris at pH 9.0 (*p < 0.001*). This suggests the critical need for testing conditions thoroughly with large-scale study designs, such as those enabled by RaPiD-mAb-MS. In this example, we weighted all modification sites equally, however, one could easily prioritize important modifications, such as those in the CDR (complementary determining region), using a weighted model. Beyond this basic machine learning use case, we envision use of RaPiD-mAb-MS data to train large-scale and more complex AI models to predict *in silico* mAb formulations, guide mAb sequence selection, PTM liability, optimum design, and bioproduction. For example, other works have described use of deep learning models (i.e., AlphaFold)^59–61^ to do intelligent mAb design; the large-scale peptide mapping data generated by RaPiD-mAb-MS could augment these existing tools to add dimensionality in mAb drug design.

### Detection of Isoaspartate via Enzymatic Labeling

While the high-throughput peptide mapping technology described above successfully identified optimal formulations, certain critical PTMs remain challenging. Specifically, the spontaneous formation of isoaspartate (isoAsp) residues through asparagine deamidation or aspartate isomerization involves species that are isobaric with their native counterparts. Consequently, direct detection via MS is impossible without chromatographic separation. To eliminate this challenge and extend the utility of RaPiD-mAb-MS to isobaric species, we next investigated selective enzymatic modification of isoAsp residues using protein L-isoaspartyl O-methyltransferase (PIMT)^62^. PIMT transfers a methyl group from S-adenosylmethionine (SAM) to the α-carboxyl group of the isoAsp side chain, generating a methyl ester that can revert to a succinimide intermediate (**Figure 7a**).

**Figure 7.**
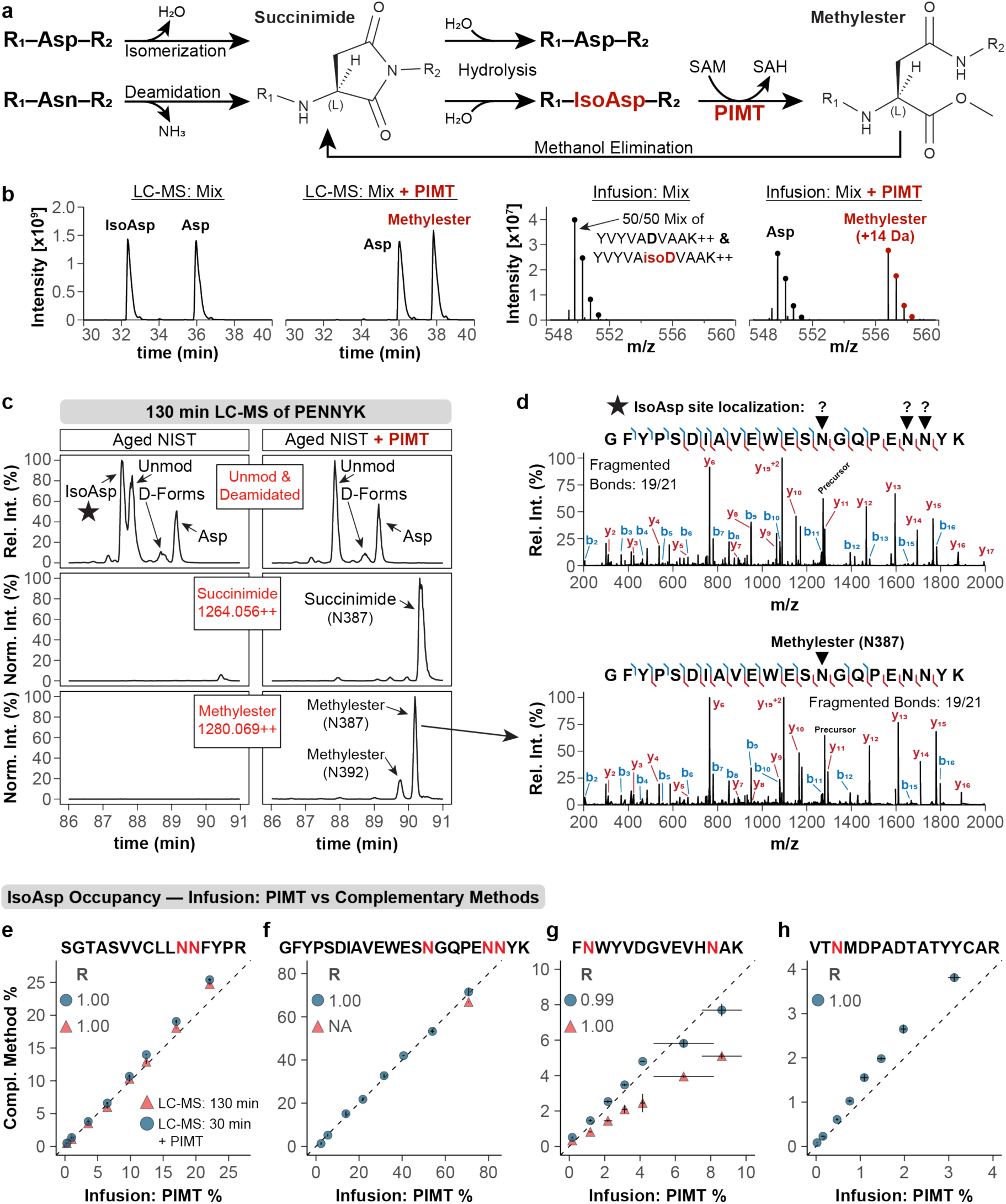
Selective labeling and detection of isoAsp modifications. PIMT transfers a methyl group from SAM to IsoAsp, generating the corresponding methylester that can revert to succinimide through methanol elimination. **b.** A 1:1 mixture of synthetic Asp and IsoAsp peptides (YVYVADVAAK and YVYVAisoDVAAK; 2+) was analyzed by nanoLC-MS (left) and direct infusion MS (right) before and after PIMT labeling. Without PIMT, Asp and IsoAsp are isobaric in MS1, whereas PIMT labeling of the IsoAsp form generates a methylester species (+14 Da), enabling separation by mass (red). **c.** Extracted ion chromatograms for the PENNYK peptide (GFYPSDIAVEWESNGQPENNYK) derived from stressed NIST mAb samples using a 130 minute nanoLC method shows partial separation of PENNYK isoAsp modified peptides. Analysis of that same sample following PIMT treatment (right) reveals conversion of the isoAsp peak and formation of succinimide and methylester species. **d.** Representative tandem mass spectra show that PIMT-modified isoAsp residues allow for simplified site localization. **e-h.** Correlation of direct infusion-based isoAsp occupancy determination for NIST mAb dilution series versus LC-MS measurement. All experiments were performed with n = 3 independent replicates. Error bars are presented as standard error of the mean (SEM).

An equimolar mixture of Asp and isoAsp-containing versions of a synthetic peptide (YVYVADVAAK) was prepared and analyzed using direct infusion and LC-MS (**Figure 7b**). Both versions of the peptide are readily observed in the LC-MS experiment but generate only a single isotopic cluster upon infusion. Analysis of the sample following incubation of these peptides with PIMT and SAM, however, presents two similar intensity isotopic clusters separated by 14 Da (**Figure 7b**). Next, we induced global isoAsp formation by stressing NIST mAb (55 °C, pH 9.3, 8 h). Even with a 130-minute LC-MS analysis, several forms of the 22 residue PENNYK peptide are only partially resolved (**Figure 7c**). Also shown is the chromatogram that results when the NIST mAb digest is treated with PIMT. PIMT labeling resulted in a complete conversion of the IsoAsp peak while Asp and D-form peaks remained, demonstrating the high efficiency and specificity of the PIMT reaction. Here we see the loss of the most prominent peak (isoAsp) and two new *m/z* peaks corresponding to the succinimide and methylester forms. Note that the resulting MS/MS spectra of these modified forms (**Figure 7d**) allow localization of the specific site of isoAspmodification – something that can be extremely challenging even by LC-MS/MS analysis when multiple possible isoAsp sites are present. Analysis of this sample using direct infusion allowed detection of isoAsp in this peptide, with occupancy estimates in excellent agreement with traditional LC-MS/MS (∼ 70% modified).

To benchmark quantitative performance, we generated a dilution curve of the stressed NIST mAb and compared the measured isoAsp levels of several peptides as measured with a traditional 130 minute LC-MS experiment (no PIMT), a short 30-minute LC-MS experiment (with PIMT), and 1 minute RaPiD-mAb-MS infusion (**Extended Data Figure 11**). **Figure 7e** presents a comparison of isoAsp measurements for the peptide having the sequence SGTASVVCLLNNFYPR – here we observe excellent linear response for the PIMT-enabled, direct infusion experiment and strong correlation with the traditional 130-minute LC-MS data. RaPiD-mAb-MS determined isoAsp occupancy values correlated strongly with the known dilution ratios (Pearson r = 0.87-0.99), demonstrating accurate quantitation. Agreement of RaPiD-mAb-MS and the traditional LC-MS approach for the other peptides was good; however, we note that in four of the seven tracked peptides we could not obtain complete isoAsp occupancy level estimates via LC-MS as the various modified versions were not fully separated. This is a known limitation for isoAsp analysis and we conclude that even in LC-MS workflows^29, 30^ the PIMT reaction method we describe here can be highly enabling.

## Discussion

Peptide mapping is a well-established and essential technique for the growing biopharmaceutical antibody therapeutic field. That said, conventional LC-MS-based peptide mapping is a time-consuming and laborious technology, such that it represents a significant throughput bottleneck in the drug development pipeline. Here we report a new technology (RaPiD-mAb-MS) that eliminates the need for chromatography, thereby enabling up to 100-fold increased data collection speed with a concomitant reduction in manual post-acquisition data analysis using RaPiD-Analyze software.

To enable RaPiD-mAb-MS we first developed and tested a plate-based sample preparation method that is simple to use, generates samples for infusion within approximately four hours, and is compatible with a wide variety of mAbs, buffers, and excipients. Reliable sample introduction is critical when omitting chromatography. The Advion NanoMate is a well-established chip-based nanoESI platform that provides quantitative performance (CVs, spray stability, and accuracy) on par with LC-MS^63, 64^. In our hands, this infusion approach proved similarly robust, maintaining signal stability across thousands of injections. Using stable isotope labeled standards we show the approach can detect modified peptides at levels less than ∼ 1% relative abundance and quantify these modifications with excellent precision and accuracy. Then, using 28 unique mAb sequences obtained from pharmaceutical companies, we obtained a median sequence coverage of 100% (mean of 97.7%) from over 1,500 experiments. Furthermore, we conducted extensive comparisons to conventional LC-MS peptide mapping across a broad spectrum of modifications (e.g., N-terminal cyclization, C-terminal clipping, deamidation, isomerization, glycosylation, and oxidation) and antibody-drug conjugates – concluding that RaPiD-mAb-MS provides highly comparable results with a fraction of the time and effort. Next, we showcase the promise and potential of RaPiD-mAb-MS with the analysis of a collection of over 1,000 mAbs that had been subjected to forced degradation. In this experiment data was collected within 1.5 days and was rapidly processed allowing for expedient selection of best performing buffers and excipients at a site-specific level. Finally, we leveraged a machine learning model to weigh the entire dataset and generate ideal conditions that would minimize all observed modifications. We imagine that with the combination of throughput offered by RaPiD-mAb-MS and use of a more advanced AI algorithm, one could predict *in silico* mAb formulation and even guide mAb sequence selection, design, and bioproduction.

Moving forward, we anticipate improvements in new peak detection (NPD). NPD is critical for identifying unexpected impurities, especially in forced degradation studies. RaPiD-mAb-MS has potential to simplify NPD as the data it produces is much simpler – i.e., eliminates the time domain component and we envision future software that can automatically flag new *m/z* features. Preliminary experiments also indicate that use of narrow MS1 *m/z* ranges (*i.e.*, selected ion monitoring) can increase the sensitivity of the method by ∼ one order of magnitude. Such capability would allow for detection of modified peptides at levels of 0.1% or lower and be especially useful for the detection of lower-level amino acid misincorporations. Additionally, early results suggest that spectral deconvolution, a promising method for determining deamidation occupancy, could be applied to RaPiD-mAb-MS using time-of-flight measurements. Spectral deconvolution works by mathematically separating overlapping native and deamidated isotope clusters thereby enabling accurate deamidation determination. This advancement would allow other mass spectrometers, even those lacking an Orbitrap, to measure deamidation values with similar speed and precision.^65–67^

Asparagine isomerization (isoAsp) is another important modification that is challenging to measure with current LC-MS technology and is not directly detectable by direct infusion. We addressed this limitation by incorporating a targeted enzymatic labeling step using PIMT. This reaction shifts the mass of isoAsp residues, allowing for their specific detection and quantification by RaPiD-mAb-MS without the need for separation. Our data confirm that this targeted workflow yields quantitative accuracy comparable to traditional LC-MS, demonstrating that even isobaric modifications can be rigorously monitored without sacrificing the throughput of the direct infusion platform. In addition, isoAsp measurements showed quantitative accuracy comparable to traditional LC-MS, with site localization benefits, demonstrating that even isobaric modifications can be rigorously monitored without sacrificing the throughput of the direct infusion platform.

By allowing for the collection of nearly 1,000 peptide maps per day, RaPiD-mAb-MS offers benefits beyond speed; it enables deep characterization at stages previously limited by analytical capacity. In early discovery, for example, deep analysis is often restricted to the final few leads. Our method allows for stress testing of dozens of candidates, ensuring selection is based on product quality and developability, not just binding affinity. Similarly, in process development, the method supports the growing trend of parallelized experimentation. It eliminates the analytical bottleneck for large-scale studies, enabling full factorial Design of Experiments that were previously impractical. Unlike conventional LC-MS, RaPiD-mAb-MS produces compact and simplified data files, facilitating the automated processing of library-scale datasets. We imagine that the accumulation of such massive, standardized data will eventually allow artificial intelligence to predict ideal formulations, antibody titer, antibody sequences, PTM liabilities, and even optimum codons.

In its present form, RaPiD-mAb-MS can detect critical modifications at an accuracy and precision commensurate with biopharmaceutical standards on widely available Orbitrap MS systems. Finally, we expect this technology to be widely applicable in multiple biotherapeutic protein modalities (e.g., fusion proteins, bi- and multi-specific antibodies, vaccines, viruses, etc.)^68–70^ and envision that the technology may be equally effective and useful across these applications.

## Methods

### Materials

Water (Optima LC/MS grade, W6-4) and methanol (Optima for HPLC, A454SK-4) were purchased from Fisher Chemical. Formic acid (85178) and Thermo Scientific™ SureSTART™ WebSeal™ Plate+ 96-Well Microtiter Plates, 220µL, V-bottom were purchased from Thermo Fisher Scientific. DL-Dithiothreitol BioUltra, ≥99.5% (43815-1G), 8M Guanidine hydrochloride solution; pH 8.5 (G7294-100ML), ammonium acetate (73594-25G-F), and Ultrafree-MC-HV Durapore PVDF 0.45um filter were purchased from Millipore Sigma. Sodium iodoacetate BioUltra, ≥99.5% (57858-5G-F) was purchased from Sigma Aldrich and Sequence Grade Trypsin, 100µg (V5117) was acquired from Promega. AcroPrep™ Advance 96-Well Filter Plates for Solvent Filtration, Pall Laboratory - 8582, AcroPrep™ Advance 96-Well Plate with 0.2 μm wwPTFE membrane, 350 µL well volume were purchased from VWR.

All 13C labeled heavy labeled peptides (Heavy Arginine (R) and Heavy Lysine(K)) were purchased from Pierce Custom Peptides with a purity >95% as lyophilized powders and “TFA removal-Acetate” selected. Heavy labeled peptides include ALPAPIEK, DTLMISR, FNWYVDGVEVHNAK, GFYPSDIAVEWESNGQPENNYK, GPSVFPLAPSSK, SFNR, VSNK, VVSVLTVLHQDWLNGK. Additionally, to look at deamidation, GFYPSDIAVEWESDGQPENNYK, GFYPSDIAVEWESNGQPEDNYK, GFYPSDIAVEWESNGQPENDYK, FDWYVDGVEVHNAK, and FNWYVDGVEVHDAK were purchased.

NIST mAb, Humanized IgG1k Monoclonal Antibody (NIST8671), SigmaMAb Universal Antibody, SigmaMAb Infliximab Monoclonal Antibody, SigmaMAb Cetuximab Monoclonal Antibody, SigmaMAb Rituximab Monoclonal Antibody were acquired from Sigma Aldrich.

### Pharmaceutical Samples

*Genentech* provided two ADCs including Genentech IgG 1 and Genentech IgG 2.

*AbbVie* provided 275 forced degraded samples in 25 mM NaPO4 pH 5.8 at 2.5mg/mL. Prior to buffer exchanging the samples’, forced degradation was performed at AbbVie in NaPO4 (pH: 5.8,7, and 8), Tris (pH: 7,8, and 9), Histidine (pH: 5.5 and 6), Acetate (pH: 5.5), and Histidine/Sucrose/PS80 (pH:6) buffers. Each of the samples had time points of 0,7,14,21,28, and 84 days as well as temperatures of 4°C, 25°C, 30°C, 40°C, and 50°C (no samples were stressed for 84 days at 50C). After forced degradation the samples were prepared using the plate method outlined below. Additionally, the forced degraded mAbs were prepared and run by LC-MS at AbbVie for comparison. The antibody−drug conjugate AB095-PZ was provided by AbbVie. The sequence of the antibody is based on human IgG1.

*Eli Lilly* provided a total of 24 unique monoclonal antibodies composed of three sets (clesrovimab, nirsevimab, and semorinemab) of the following Fc mutations per molecule: IgG1, IgG1+YTE, IgG1-AAS, IgG1-AAS+YTE, IgG2, IgG2+YTE, IgG4, and IgG4+YTE. The samples were provided in 1X PBS buffer and normalized to 2.2mg/mL. Following concentration normalization, 50ug of each molecule was stressed for 0,7,14, and 28 days at 50°C. After forced degradation the samples were prepared using the plate method outlined.

*Johnson & Johnson*: Forced degraded samples were provided for four different mAbs including NIST mAb, Belimumab, Nivolumab, and MAB1. For each molecule, there were three variables (control- no temperature, thermal stress (40 °C for 14 days), and photostability (100 W/m^2^ for 3 hrs (30 Whrs))) with a myriad of conditions for each plate with both “buffer” and “oxidation” matrices being used. Refer to **Supplemental Table 1-3** for the full outline of the samples.

### Antibody Sample Plate Method Preparation

*Protein Denaturation, Reduction, and Alkylation*: To an AcroPrep™ 96 well plate, 50 µL 8M GnHCl was pipetted into each well followed by 50 µg of antibody sample (<20 µL). To this, 2.5 µL of 0.5M DTT was added and incubated at 37°C for 15 minutes. After incubation 3 µL 1M IAA was pipetted followed by incubating at room temperature in the dark for 15 minutes. Following incubation, each well was quenched by adding 3.6 µL of the 0.5M DTT solution. *Protein Precipitation and Washing*: 200 µL MeOH was added to each well and mixed thoroughly by pipetting up and down while limiting any bubble formation to precipitate the protein. This was then incubated at room temperature for 10 minutes. After incubation, the AcroPrep™ plate was placed on a 96 deep-well format plate and centrifuged at 4,000 RPM (2272g) for 10-15 minutes with the filtrate being discarded. 200µL of 80% methanol in water was added to each well and centrifuged at 4,000 RPM (2272g) for 10-15 minutes with the filtrate being discarded. The washing was repeated four more times (5X total) discarding the filtrate after each round of centrifugation. Before moving on to the next step we confirmed that there was very little or no liquid remaining in the wells. If there was still significant liquid, centrifugation was done until all wells are empty (wells should not look physically dry). *Protein Digestion and Final Elution*: To each well 90 µL of 100mM ammonium acetate solution was added followed by adding 10 µL of trypsin (e.g. 2µg total per well). After a 2-hour incubation at 37°C, the digestion was quenched by adding 10 µL of 10% formic acid solution. A new clean 96-well plate was placed under the AcroPrep™ plate and centrifuged for 10 minutes at 4,000 RPM (2272g) to collect peptides in the filtrate. The samples then had a 96-well plate foil applied. These samples can be used directly for MS direct infusion analysis.

### Direct Infusion Mass Spectrometry

Experiments were performed on Thermo Scientific Orbitrap Eclipse Tribrid and Orbitrap Astral MS systems. Due to the NanoMate supplying the voltage for ionization, the MS ionization voltage was set to zero with a capillary temperature of 90°C.

Orbitrap Eclipse Tribrid: MS1 experiments were conducted in the Orbitrap with a resolving power of 480K and a scan range of 200-2000m/z. The RF lens was set to 30% with a normalized AGC target of 250% (1E6 ions). Three microscans were applied with data being collected in profile mode with a positive polarity. Targeted MS2 Orbitrap HCD scans were achieved with an isolation window of 7m/z with a normalized collision energy of 30%. The Orbitrap resolution was set to 480K with a scan range of 200-2000m/z and a RF lens set to 50%. Normalized AGC target was set at 400% (2E5 ions) with a custom maximum injection time of 400ms. One microscan was employed with data being acquired in positive profile mode with targets looping in an unscheduled time mode. For DDA MS/MS, the same MS1 master scan settings were used for target selection. Extensive filtering was used to select the DDA targets. First, a relative intensity of 0.5% was applied followed by charge state 1-6 and MIPS designation of “peptide” with the most abundant peak being isolated. Both assigned and unassigned isotope exclusion were performed with a dynamic exclusion of 60 seconds. DDA Orbitrap HCD isolation was achieved with the quadrupole and an isolation window of 1m/z. HCD collision energy of 30% was used at a resolution of 15K and an AGC target of 500% (2.5E5) with a maximum injection of 150ms and data type of centroid. Orbitrap Astral: MS1 experiments were conducted in the

Orbitrap with a resolving power of 480K and a scan range of 200-2000m/z. The RF lens was set to 50% with a normalized AGC target of 250% (2.5E6 ions) and a max inject time of 50ms. Three microscans were applied with data being collected in profile mode with a positive polarity. Targeted MS/MS Orbitrap HCD scans were achieved with an isolation window of 4m/z with a normalized collision energy of 30%. The Orbitrap resolution was set to 480K with a scan range of 200-2000m/z and an RF lens set to 50%. Normalized AGC target was set at 400% (4E5 ions) with a custom maximum injection time of 100ms. One microscan was employed with data being acquired in positive profile mode with targets looping in an unscheduled time mode. For DDA MS/MS, the same MS1 master scan settings were used for target selection with an isolation window of 1m/z and normalized collision energy of 30% for HCD. The detector used was the Astral with a scan range of 200-2000, a standard AGC target, maximum injection time of 10ms, 1 microscan, and a datatype of Centroid. **Supplementary Table 4** summarizes the MS instruments used for each direct-infusion MS and LC-MS experiment, organized by figure.

### TriVersa NanoMate

All direct infusion experiments were conducted with an Advion TriVersa NanoMate. For delivery of samples on the NanoMate system, pre-piercing with a mandrel was performed along with venting of the headspace prior to infusion. Additionally, an air gap prior to sample engaging the chip was employed with an aspiration delay and equalization delay of 2 seconds and a volume after delivery of 0.5uL. A positive voltage of 1.65-1.75kV was applied to each sample with a voltage timing of “Voltage Before” and a delay of zero seconds as well as 1 psi of nitrogen gas pressure. For large studies, contact closure was achieved using the NanoMate system with an output signal being sent for 2.5 seconds on Rel1. Sample volume and delivery time were variable based on experiment with volumes between 1-3uL being picked up and infusion times between 30-120 seconds.

### LC-MS

Sample analysis was performed on an Acquity UPLC CSH C18 column held at 50°C (100 mm x 2.1 mm x 1.7 μm particle size; Waters). 10-15 μL of sample was injected by a Vanquish Split Sampler HT autosampler (Thermo Scientific). Separations were performed using a Vanquish Binary Pump (0.5 mL/min flow rate; Thermo Scientific) with the following gradient: initial conditions of 100% Mobile phase A (0.1% formic acid in water) for 1 minute, then linear increase to 35% Mobile phase B (0.1% formic acid in acetonitrile) over the next 38 minutes. Over 1-minute, Mobile phase B was ramped to 100% and was maintained for 4 minutes before returning to 0% mobile phase B over the next 1 minute and equilibrating at 0% for a remaining 5 minutes. The LC system was coupled to an Orbitrap Tribrid Eclipse or Ascend through a heated electrospray ionization (HESI) source (Thermo Scientific). Source conditions were as follows: HESI and capillary temperature at 350°C, vaporization temperature at 300°C, sheath gas flow rate at 50 units, aux gas flow rate at 15 units, sweep gas flow rate at 2 units and, spray voltage at |3.5 kV| in positive mode. The MS performed full MS and MS/MS data collection in positive mode. Acquisition parameters for full MS scans were 120,000 resolution, 4 × 10e5 automatic gain control (AGC) target, 250 ms ion accumulation time (max IT), and 350 to 2000 m/z scan range with RF lens at 30%. MS2 scans were performed at 30,000 resolution, 5 × 10e4 AGC target, 60 ms max IT, 1.6 m/z isolation window, and HCD at 30%.

### AbbVie Data Collection

Automated peptide digest was performed on a Hamilton Starlet, with all steps taking place at room temperature. Briefly, proteins were denatured with 4 M guanidine, followed by reduction with 20 mM DTT for 60 minutes and alkylation with 40 mM iodoacetic acid for 60 minutes both at room temperature (pH 8), after which proteins were desalted with 1 mL PHYTIPS filled with 600 μL 10K resin (Phynexus). Digestion was performed with Trypsin/Lys-C enzyme (Promega) at a 1:25 enzyme:substrate ratio for 2 hours in digestion buffer (50 mM Tris-HCl, 1 M Urea, 20 mM L-methionine, pH 7.5). Digest was halted by addition of TFA to 0.5% final concentration. Samples were injected to a Thermo Fisher Scientific Vanquish-Fusion LUMOS LC-MS system with the following column: Waters ACQUITY UPLC BEH 300; C18; 1.7 μm Dimension: 2.1 mm × 150 mm, PN:186003687. Mobile Phase A: LC/MS water with 0.08% FA/0.02% TFA; Mobile Phase B: LC/MS acetonitrile with 0.08% FA/0.02% TFA. A multi-step gradient from 1 to 40% MPA to MPB was performed, with a top-5 MS/MS setting (DDA, MS1 60,000 resolution, scan range 250-2000 m/z; MS2 15,000 resolution). Data was analyzed in Protein Metrics Byos (PMI by Dotmatics). After the initial peptide search, in silico match to existing was performed, and relative quantification estimated by XIC ratio.

### Johnson & Johnson Sample Preparation

Methods and Material for High throughput Screen (HTS). *Reagents and Materials.* NIST mAb and humanized IgG (MAB1) were manufactured at J&J Innovative Medicine and formulated in a histidine -based buffer. Belimumab (Benlysta) and Nivolumab (Opdivo) were purchased from Myonex (Norristown, PA) and diluted using high purity water. L-histidine was purchased from Millipore Sigma (Burlington, MA). Sucrose was purchased from Pfanstiehl (Waukegan, IL), citric acid monohydrate, sodium citrate dihydrate, sodium acetate trihydrate, sodium chloride (NaCl), sorbitol, L-histidine monohydrochloride monohydrate, tris(hydroxylmethyl) aminomethane, L-proline, L-methionine, L-arginine monohydrochloride, iron (III) chloride (FeCl_3_) hexahydrate, and hydrogen peroxide (H_2_O_2_) solution were purchased from Sigma-Aldrich (St. Louis, MO). Acetic acid glacial and EDTA were purchased from VWR (West Chester, PA). Hydrochloric Acid was purchased from JT Baker (Avantor, Radnor, PA)

#### Automated Liquid Handler

96 plate-based formulations were prepared using a MICROLAB STAR (Hamilton, Reno, NV)) liquid handler, as outlined in **Supplementary Tables 1-3**. The liquid handler can be programmed to pipette and mix the various solutions in the order desired. To set up the buffer/pH/excipient plate, concentrated stock solutions were prepared for protein, buffer (citrate, histidine, acetate, tris) and excipients (none, proline, arginine, NaCl, sucrose, sorbitol). The molarity and pH of target formulations were achieved with calculated concentrations of conjugate acid and conjugate base, whereas relevant target excipient concentrations were attained using serial dilutions. The final target volume in each well was achieved by adding high purity water. The oxidation plate was prepared by mixing excipients as outlined in Supplementary Table 2 in microliter volumes. The well volumes for the “mother” plate ranged from 300 µL to 500 µL. “Daughter” plates were then generated by aliquoting material from the mother plate into 3 separate plates, sealed, and stored at each of the appropriate stress conditions.

#### Protein Concentration Measurements

Protein concentration of the control plate was measured based on the absorbance at 280 nm. Samples were tested neat at 3 μL volume in a microfluidic plate using a Lunatic UV-Vis spectrometer (Unchained Labs, Pleasanton, CA).

#### High Temp and Photo Exposure

A total of twelve (4 proteins X 3 daughter plates) 96-well plates were prepared for the buffer, pH, excipient and oxidation studies. Each protein plate was designated for the control, high temp, or photo exposure condition. 96-well plates for the control and high temp conditions were sealed using silicone film and aluminum adhesive film to protect the samples from evaporation. Control samples were frozen at -40°C immediately after preparation. Plates exposed to high temp stress were stored at 40°C for 2 weeks using a controlled chamber (Thermofisher, Waltham, MA). The photo exposure study was performed by placing the appropriate plates without any cover into a photostability chamber (Environment Specialties, Raleigh, NC) with the UV lamp set to 10 watts/m^2^ for 3 hours, yielding 30 watt-hours of exposure. The light-exposed samples were then sealed using silicone and aluminum film. High temp and photo-stressed samples were stored at -40°C and all samples were shipped under frozen conditions for analysis.

### Software

RaPiD-Analyze software was written in C# to automate data analysis. Proteins are digested in silico. For each charge state of each putative peptide within the mass range, the theoretical isotopic distribution is computed. The abundance within the mass tolerance of each theoretical isotopic peak above a threshold is extracted from the averaged MS1 spectrum. The scale factor is determined which minimizes the sum of the squared differences between the experimental and theoretical isotopic abundances. Total abundance is calculated by summing the scale factor-weighted theoretical isotopic abundance over all isotopic peaks. An abundance-weighted root mean square isotopic error is also calculated for each isotopic cluster, and those with an isotopic error over a threshold are deemed as not detected.

The occupancy of each isoform is then calculated as the abundance of the isoform of interest divided by the sum of the abundance of all isoforms of the same peptide. Only peptides without missed cleavages are considered. Only charge states detected for all isoforms are used except when this is not possible, in which case all charge states are used. Abundances are normalized by charge state.

For deamidation occupancy by MS2, all singly charged fragment ions are evaluated to determine if they both contain at least one potential deamidation sites and are free from interference from other fragment ions. The abundance of both unmodified and modified fragment ions are then determined similarly to MS1 by generating theoretical isotopic distributions and optimizing for the scale factor which minimizes the sum of squared differences between experimental and theoretical isotopic abundances. We then set up a series of simultaneous linear equations whereby the occupancies of each possible deamidation site are unknowns and the observed occupancies as determined by ratio of modified over total fragment ions containing those sites are knowns. These equations are also weighted by the total abundance of fragment ions contributing to the occupancy. This yields an overdetermined system of equations for which the optimal solution can be determined by least squares.

### Oxidation Forced Degradation Titration

NIST mAb, Infliximab, Cetuximab, Rituximab, and Sigma mAb Universal antibody were prepared at ∼10mg/mL in 25mM ammonium acetate buffer (excluding NIST mAb which is already in solution when purchased). From each antibody, 18uL were taken and mixed with 110uL of 0.025% H2O2 in 8M GnHCl and was oxidized for 90 minutes. Each sample was then split and precipitated in 500uL of methanol in an Ultrafree-MC-HV Durapore PVDF 0.45um filter tube. This was allowed to sit for 5 minutes and then was centrifuged at 10,000g for 1 minute. The samples were rinsed with 500uL of 90% methanol four times. 91uL of GnHCl buffer was added to each sample which was vortexed for ∼1 minute followed by being centrifuged for 1 minute to obtain the oxidized antibody. The two samples were combined and found to be at a ∼1mg/mL force degraded antibody. From each of the bulk native antibody samples that had not been oxidized, a solution of each antibody was prepared at 2.5mg/mL. A titration of the native antibody (2.5mg/mL) and the oxidized antibody (1mg/mL) was performed in the following ratios (stressed µL)/(native µL): (50/0, 31.3/7.5, 12.5/15, 6.3/17.5, 3.1/28.1, 0/20) and then prepared using the antibody plate digestion method with an initial volume in the well of 10uL of 8M GnHCl instead of 50uL. Each sample was run in triplicate on both LC-MS and NanoMate direct infusion with a Tribrid MS.

### ^13^C Peptide Titration-LOD

1mg of dried lyophilized powder for each heavy peptide was obtained and dissolved in 1mL of solvent to obtain a solution of 1mg/mL of each heavy peptide. Once each dissolution was achieved a calculated volume of each heavy peptide solution was pipetted into a single tube to achieve 50pmol/µL total of each ^13^C peptide with the remaining volume being brought up to 1mL total with water. Once the 50pmol/µL for each ^13^C peptide was achieved in one solution, this was then serial diluted from 50pmol/µL to 8fmol/µL (16.7µL of the initial 50pmol mixture was taken and pipetted into 33.3µL of ammonium acetate buffer and mixed thoroughly with an additional 8 dilutions performed). From this, 10µL of each ^13^C serial dilution solution was taken and spiked into 90µL of NIST mAb peptides (started at 0.32mg/mL) that had been previously prepared using the plate method. This provided samples from 5000 to 0.76 fmol/µL of ^13^C peptides in a constant background of NIST mAb. The samples were then run in triplicate using LC-MS and on NanoMate direct infusion with a Tribrid MS.

### ^13^C Peptide Titration-Deamidation

Both ^13^C native (FNWYVDGVEVHNAK, GFYPSDIAVEWESNGQPENNYK -1mg each) and deamidated (GFYPSDIAVEWESDGQPENNYK, GFYPSDIAVEWESNGQPEDNYK, GFYPSDIAVEWESNGQPENDYK, FDWYVDGVEVHNAK, and FNWYVDGVEVHDAK-0.083mg each) peptides were obtained as dried lyophilized powders and dissolved in 1000 and 325uL of solvent respectively. Once dissolved, the amount of volume of each heavy peptide was determined to achieve 100%-0% deamidation, decreasing deamidation by a factor of 2 each dilution, with the final pmol amount of each isoform being 50pmol total. The starting sample, 100% deamidation, had the following: 0pmol GFYPSDIAVEWESNGQPENNYK, 25pmol GFYPSDIAVEWESDGQPENNYK, 18.8pmol GFYPSDIAVEWESNGQPEDNYK, and 6.3pmol GFYPSDIAVEWESNGQPENDYK. Additionally, there was 0pol FNWYVDGVEVHNAK, 16.7pmol FDWYVDGVEVHNAK, and 33.3pmol FNWYVDGVEVHDAK. To make the remaining 8 samples, the pmol amounts of each deamidated heavy peptide were halved while the amount of native peptide was increased to achieve the 50pmol of the overall isoform of each peptide. From each of these samples with varying amount of deamidated and native peptides, 10uL was taken and diluted in 90uL of NIST mAb peptides (started at 0.32mg/mL) that had been previously prepared using the plate method. The samples were then run in singlicate on the LC-MS and in triplicate with NanoMate direct infusion on a Tribrid MS.

### Linear model determination

Linear model for **Extended Data Figure 10** was calculated as follows: percent modification was calculated from 16 modification sites across 14 unique peptide sequences on NIST mAb, across 272 unique combinations of stressor, buffer, pH and excipient. Method stastmodels.formula.api.mixedlm from the Statsmodels library (version 0.14.0) in Python was used to calculate the model, specified as: modification_percent ∼ stressor + excipient + buffer_pH_combo, with random effects given with groups=’peptide’.

### PIMT-specific reagents

Guanidine hydrochloride (G4505), glycine (G8898), and sodium hydroxide (221465) were purchased from Sigma-Aldrich. 1 M bicine buffer, pH 8.5 (40121184-1), was purchased from Bioworld.

### PIMT labeling of synthetic peptide standards

Pierce custom peptides (Asp: YVYVADVAAK; IsoAsp: YVYVAisoDVAAK, where isoD denotes isoaspartate at the Asp position) were mixed 1:1 to a final concentration of 6.66 µM total (3.33 µM each) in 10 mM ammonium acetate containing 25% methanol (v/v) and 200 µM S-adenosylmethionine (SAM), adjusted to pH 5.7, in a total volume of 75 µL. Recombinant PIMT was supplied via a custom production order (Promega) and added at 0.8 µg per reaction. Reactions were incubated overnight at 37 °C and quenched by adjusting to 0.5% (v/v) formic acid prior to direct infusion MS analysis.

### NIST mAb aging (stress) treatment for PIMT labeling

The aging buffer was prepared by dissolving guanidine hydrochloride (GnHCl) to 6 M and adjusting to 50 mM glycine using 1 M glycine-NaOH (pH 9.8), yielding a final buffer of 5.7 M GnHCl, 50 mM glycine, pH 9.8. To generate “Aged NIST,” 5 µL NIST mAb (50 µg) was mixed with 42.5 µL aging buffer (47.5 µL total), yielding a final pH of 9.3, and incubated at 55 °C for 8 h. Next, 7.5 µL of 1 M bicine (pH 8.5) was added to a final volume of 55 µL, readjusting the pH to 8.5. For mock-treated (native) controls, 5 µL NIST mAb (50 µg) was combined with 42.5 µL 5.7 M GnHCl (without glycine; pH unadjusted), omitting the 55 °C aging step. Then, 7.5 µL of 1 M bicine (pH 8.5) was added (55 µL total) to adjust the pH to 8.5.

### PIMT labeling of NIST mAb

Following the NIST mAb aging (stress) treatment, Native NIST mAb and Aged NIST mAb were mixed to generate a nine-point native-to-aged dilution series (Samples 1-9; mixing ratios in Extended Data Figure 11b). Samples (55 µL per well) were loaded onto an AcroPrep™ 96-well plate and processed using the Antibody Sample Plate Method Preparation workflow (denaturation, reduction/alkylation, precipitation/washing, and on-plate tryptic digestion) as described above. The only deviation was that tryptic digestion (2 h at 37 °C) was carried out in 110 µL of 25 mM ammonium acetate (rather than 90 µL of 100 mM ammonium acetate). Following digestion, samples were not acidified (i.e., the formic acid quench was omitted). Instead, peptides were eluted from the filter plate in the digestion buffer. Samples were split into two aliquots (50 µL each) for analysis with or without PIMT labeling. For the mock-treated (-PIMT) aliquot, formic acid was added to 0.5% (v/v), and samples were stored at -40 °C until DI-MS analysis. For the PIMT-treated (+PIMT) aliquot, PMSF was added to 1 mM and samples were incubated for 1 h at 37 °C to inactivate co-eluted trypsin. Samples were then adjusted to pH 5.7 by addition of formic acid, supplemented to 200 µM S-adenosylmethionine (SAM), and recombinant PIMT was added (1 µg per reaction). Reactions were incubated overnight at 37 °C and quenched by addition of formic acid to 0.5% (v/v) prior to DI-MS analysis.

### LC-MS analysis of PIMT experiments

For nanoLC-MS analysis of PIMT-related experiments, a Vanquish Neo UHPLC system (Thermo Fisher Scientific) was coupled to an Orbitrap Ascend (custom peptide PIMT labeling; Fig. 7b), an Orbitrap Exploris 240 (PIMT-labeled NIST mAb-derived peptides), or an Orbitrap Astral mass spectrometer (parallel reaction monitoring, PRM, of PIMT-labeled NIST mAb-derived peptides) (all Thermo Fisher Scientific). All instruments were equipped with a Nanospray Flex ion source. Peptides were separated on a 40 cm in-house packed fused-silica column (C18, 1.7 µm, 130 Å) maintained at 50 °C using a custom-built column heater. The spray voltage was set to 2.0 kV, and the ion-transfer-tube temperature was 275 °C (Ascend) or 280 °C (Exploris 240 and Astral). For custom peptide experiments, 3 pmol total peptide (1.5 pmol of each synthetic peptide in a 1:1 mix) was injected and separated using a 73 min active gradient (84 min total). For NIST mAb peptide experiments, 2 pmol per peptide was injected and separated using either a short gradient (20 min active; 31 min total) or a long gradient (120 min active; 131 min total). Separations were performed at 300 nL/min using an active gradient from 5% to 46% solvent B (solvent A: 0.2% formic acid in water; solvent B: 80% acetonitrile with 0.2% formic acid).

PIMT experiments on custom peptides were analyzed on an Orbitrap Ascend using an MS1-only method. Full MS1 scans were acquired in the Orbitrap at a resolution of 240,000 over an m/z range of 370–2000 with a maximum injection time of 100 ms, a normalized AGC target of 250%, RF lens setting of 40%, and 3 microscans.

PIMT-labeled NIST mAb peptides were analyzed on an Orbitrap Exploris 240 using a data-dependent acquisition (DDA) approach: full MS1 scans were acquired in the Orbitrap at a resolution of 240,000 over an m/z range of 350-1350 with a maximum injection time of 100 ms, normalized AGC target of 500%, RF lens setting of 40%, and a single microscan. MS1-to-MS2 precursor selection (applied scan filters) used monoisotopic precursor selection set to Peptide, charge-state screening of 1-6, and dynamic exclusion (20 s exclusion duration; ±5 ppm mass tolerance). MS2 spectra were acquired using HCD with a normalized collision energy of 25%, an isolation window of 0.9 m/z, and Orbitrap detection at a resolution of 15,000 with a maximum injection time of 22 ms, normalized AGC target of 250%, RF lens setting of 70%, and a single microscan.

On the Orbitrap Astral, PIMT-labeled NIST mAb peptides were analyzed using an MS1–PRM method. MS1 settings were matched to the Exploris 240 method except that the normalized AGC target was set to 250% and the maximum injection time was set to 50 ms. For PRM acquisition, an inclusion list was generated for the monoisotopic precursor m/z values of each target peptide and for the following variants: unmodified, deamidated, succinimide, and methylester. PRM scans were acquired with an isolation window of 0.8 m/z using HCD at a normalized collision energy of 26%, a scan range of m/z 200–2000, a maximum injection time of 20 ms, a normalized AGC target of 500%, RF lens setting of 40%, a single microscan and an unscheduled time mode.

### DI-MS analysis of PIMT experiments

All PIMT-related DI-MS experiments were performed using an Advion TriVersa NanoMate coupled to an Orbitrap Ascend.

Custom peptide experiments were analyzed using an MS1+SIM method design. Method duration was set to 1 min and ion transfer tube temperature to 200 °C. Full MS1 scans were acquired in the Orbitrap at a resolution of 480,000 over an m/z range of 370-2000 with a maximum injection time of 100 ms, a normalized AGC target of 250%, RF lens setting of 40%, and 3 microscans. SIM scans were acquired at a resolution of 240,000 over an m/z range of 350-2000 with a maximum injection time of 507 ms, a normalized AGC target of 250%, RF lens setting of 40%, and a single microscan. All YVYVADVAAK/YVYVAisoDVAAK-derived peptide species were targeted in SIM mode using a 23 m/z isolation window in unscheduled mode.

PIMT-labeled NIST mAb peptides were analyzed using the same method structure described in the general section “Direct Infusion Mass Spectrometry.” The total method duration was 2 min and the ion transfer tube temperature was set to 275 °C. Briefly, from 0.0-1.4 min, full MS1 scans were alternated with unscheduled targeted MS2 (tMS2) scans directed at N-containing NIST mAb peptides; from 1.4-2.0 min, a DDA segment was included. MS1 scans were acquired in the Orbitrap at a resolution of 480,000 over an m/z range of 200-2000 with a maximum injection time of 100 ms, a normalized AGC target of 250%, RF lens setting of 50%, and in 3 microscans. tMS2 scans were acquired at a resolution of 480,000 (scan range mode: Auto) with RF lens set to 50%, a normalized AGC target of 1000%, a maximum injection time of 500 ms, and HCD at a normalized collision energy of 26%; isolation windows were 5.0 m/z for doubly charged and 4.3 m/z for triply charged precursors. In the DDA segment (ddMS2), precursors were isolated with a 1.0 m/z window and fragmented by HCD (normalized collision energy 26%), and MS2 spectra were acquired in the Orbitrap at a resolution of 15,000 over an m/z range of 150-2000 with a maximum injection time of 27 ms, a normalized AGC target of 500%, and a single microscan. These tMS2-containing runs were used for the −PIMT subset of the experiment. For the +PIMT subset, the method was identical except that tMS2 scans were replaced by multiplexed SIM scans (multiplex ions enabled) acquired at a resolution of 480,000 over an m/z range of 350-2000 with a maximum injection time of 300 ms, a normalized AGC target of 500%, RF lens setting of 50%, a single microscan, and unscheduled time mode. Multiplexing groups were designed such that the succinimide, unmodified, and methylester forms of a given peptide were combined into a single scan; individual targets were isolated using 6.0 m/z (z = 1), 4.5 m/z (z = 2), or 4.0 m/z (z = 3) isolation windows.

### IsoAsp Occupancy Calculations

IsoAsp occupancy was calculated from monoisotopic MS1 signals, using peak intensities for DI-MS and chromatographic peak areas (AUC) for LC-MS. For DI-MS, monoisotopic intensities were extracted from averaged multiplexed SIM spectra targeting the unmodified, succinimide, and methylester species using a 6 ppm mass tolerance. Only the most abundant charge state per peptide was targeted (i.e., no cross-charge summation). SIM scans were chosen as an MS1-level equivalent of the targeted MS2 (tMS2) scans applied for general deamidation assessment. An analogous single-charge-state quantitation strategy was applied to LC-MS data (MS1 AUCs in Skyline; 6 ppm) to enable direct comparison across platforms. For PIMT-based measurements, IsoAsp levels were quantified as the sum of the succinimide and methylester signals, and IsoAsp occupancy (%) was calculated as:

100 × (I_succinimide + I_methylester) / (I_unmodified + I_succinimide + I_methylester),

where ”I” denotes the monoisotopic MS1 signal (peak intensity for DI-MS or AUC for LC-MS). For LC-MS measurements without PIMT, IsoAsp occupancy was calculated directly as:

100 × IsoAsp / (IsoAsp + unmodified).

Because succinimide can also be generated as a processing artifact and methylester signals may include low-level background or interfering peaks, IsoAsp occupancy values were normalized to the occupancy value of Sample 1 (S1, 100% Native) of the NIST mAb dilution series (Extended Data Fig. 11b). Consequently, S1 was used as the normalization reference and was omitted from correlation analyses.

## Acknowledgments

We are grateful for NIH grants P41 GM108538, R35 GM118110, and R35 GM126914 for partial financial support of this work. We also thank the University of Wisconsin-Madison and the Morgridge Institute for Research. We thank Wendy Sandoval and Wilson Phung of Genentech for providing mAb samples and helpful discussions. We thank Johnson & Johnson for sourcing clinical grade commercial antibodies and manufacturing the NIST mAb antibody for this study.

## Code Availability

The custom software suite used for data analysis is available through CeleramAb, Inc. (www.celeramab.com).

## Contributions

A.S.H., A.Z.S., M.M., L.R.S., J.J.C, and K.L.M. contributed to 96 well-plate and MS method development.

A.Z.S., M.M., J.S.C., A.S.H., and B.R.P. performed mass spectrometry sample preparation and/or data acquisition.

A.Z.S., M.M., J.S.C., and B.J.A. performed mass spectrometry data processing.

C.D.W., P.S., J.J.C., A.Z.S., I.J.M., and E.M.G. conceived and developed data processing software.

A.Z.S., M.M., A.S.H., P.S., J.S.C., B.J.A., and J.J.C. contributed to the figure content and design.

A.M.G.T., M.P.G., J.A., M.B., B.R.P., G.L., and H.P.G. provided samples, contributed to experiment design, and expert analysis.

A.Z.S., M.M., B.J.A., and J.J.C. wrote the manuscript. A.Z.S., M.M., L.M.S. and J.J.C. edited the manuscript

