## Extended and Supplementary Data for "Rapid Peptide Mapping of Monoclonal Antibodies with Direct Infusion Mass Spectrometry"

#### Extended Data Figures

Extended Data Figure 1: RaPiD-mAb-MS $^{13}$C peptide sensitivity and sequence coverage across samples

Extended Data Figure 2: Comparison of RaPiD-mAb-MS to LC-MS peptide mapping

Extended Data Figure 3: Quantifying deamidation by conventional LC-MS and RaPiD-mAb-MS

Extended Data Figure 4: AbbVie antibody-drug conjugate (AB095-PZ) analysis by MS1 and DDA MS/MS: THTCPPCPAPELLGGPSVFLFPPKPK peptide

Extended Data Figure 5: AbbVie antibody-drug conjugate (AB095-PZ) analysis by MS1 and DDA MS2: Small Peptides.

Extended Data Figure 6: Global RaPiD-mAb-MS and LC-MS peptide mapping comparison for NIST mAb.

Extended Data Figure 7: RaPiD-mAb-MS and LC-MS peptide mapping comparison of other modification sites.

Extended Data Figure 8: Heatmap for oxidation conditions.

Extended Data Figure 9: Bar plots of NIST mAb peptides from oxidation conditions with single methionine and tryptophan present as well as the combination in one peptide.

Extended Data Figure 10: Heatmap for NIST mAb for both buffer and oxidation conditions and effect of tested stressors, excipients, buffer and pH on average modification percent on 16 modification sites in NIST mAb.

Extended Data Figure 11: Accurate isoAsp quantitation in stressed NIST mAb by DI-MS and LC-MS, benchmarked across a native-to-aged dilution series.

### Extended data

#### Extended Data Figure 1: RaPiD-mAb-MS $^{13}C$ peptide sensitivity and sequence coverage across samples.

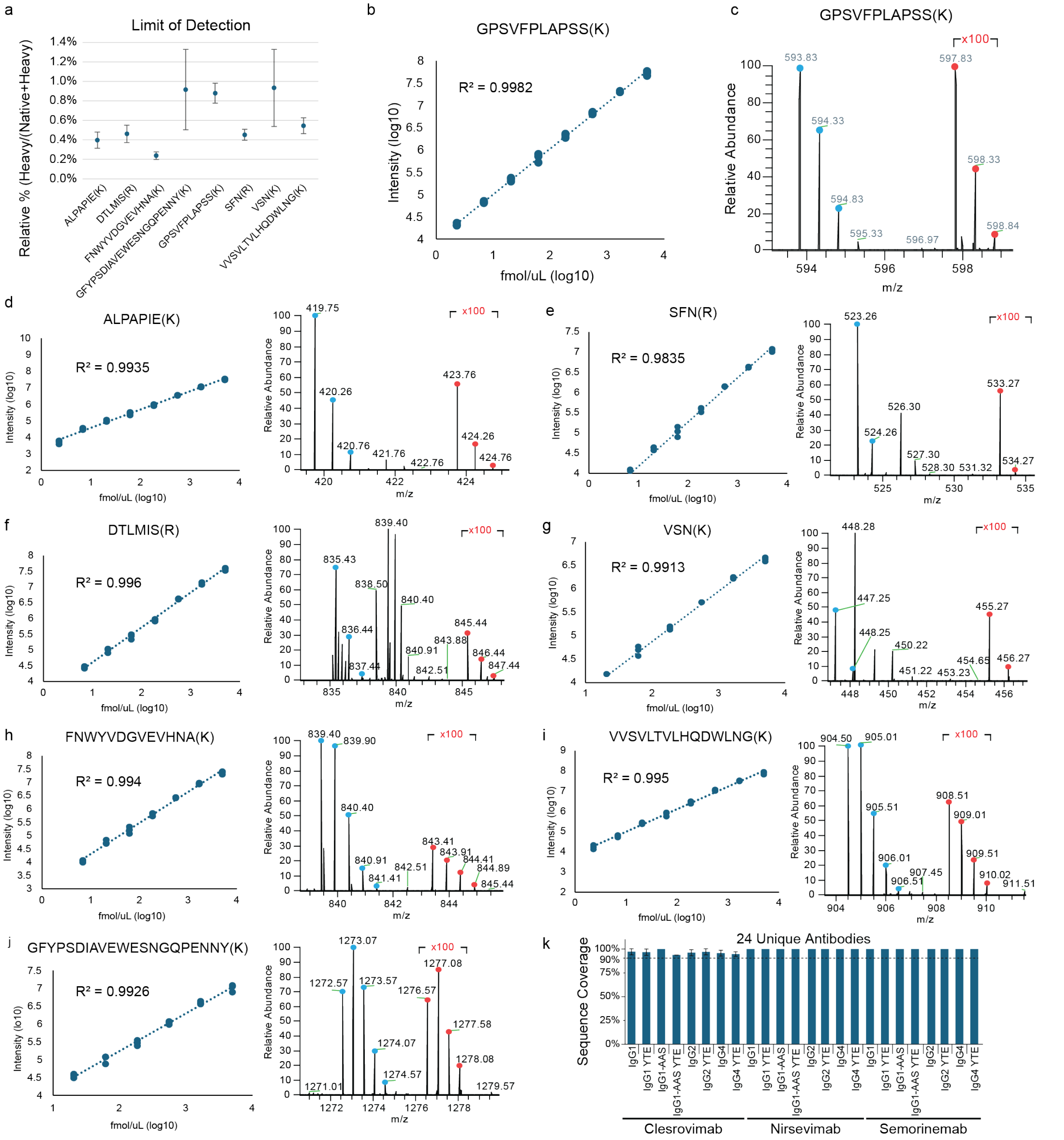


**a**, Limit of detection calculated for 8 heavy peptides (based on 3X spectral signal-to-noise, n=3) spiked into the background of NIST mAb peptides showing high sensitivity down to ~1% when compared to the native peptide. As an example, the log10 titration curve is shown (**b**) for the GPSVFPLAP(K) $^{13}C$ peptide along with a zoom in of the $^{13}C$ peptide by 100x (red) compared to its native peak (blue) (**c**). The remaining peptides are shown in **d-i**. **j**, sequence coverage of 24 unique antibodies (n=3). **k**, Sequence coverage for 24 antibodies derived from the Clesrovimab, Nirsevimab, and Semorinemab base sequences.

**Extended Data Figure 2: Comparison of RaPiD-mAb-MS to LC-MS peptide mapping.**

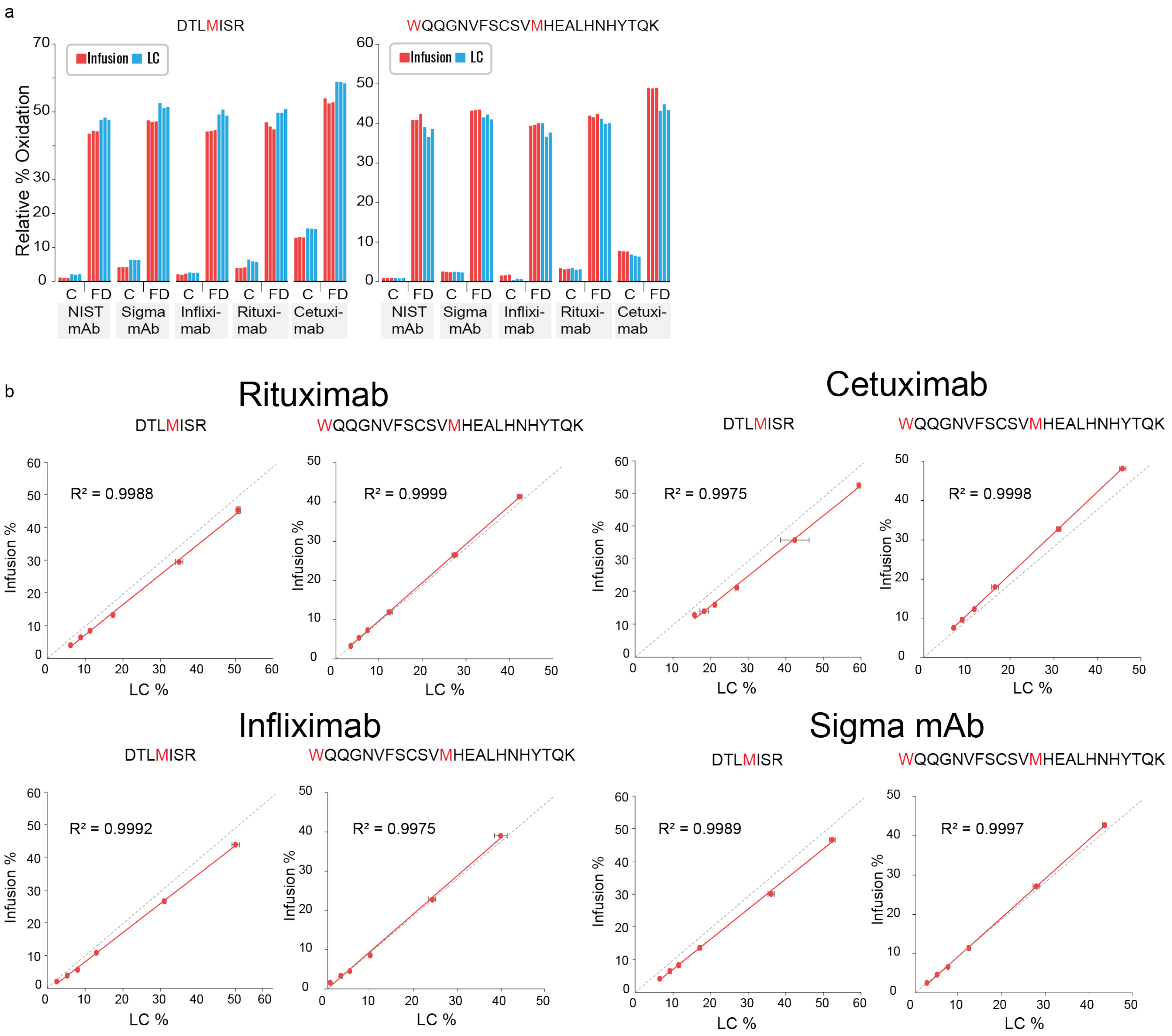


**a,** Single timepoint comparison and **b,** full plots of RaPiD-mAb-MS and LC-MS quantification measurements for oxidation of M containing peptides across five mAbs (n=3, dashed line is y=x). Abbreviations: C, control; FD, forced degraded.

**Extended Data Figure 3: Quantifying deamidation by conventional LC-MS and RaPiD-mAb-MS.**

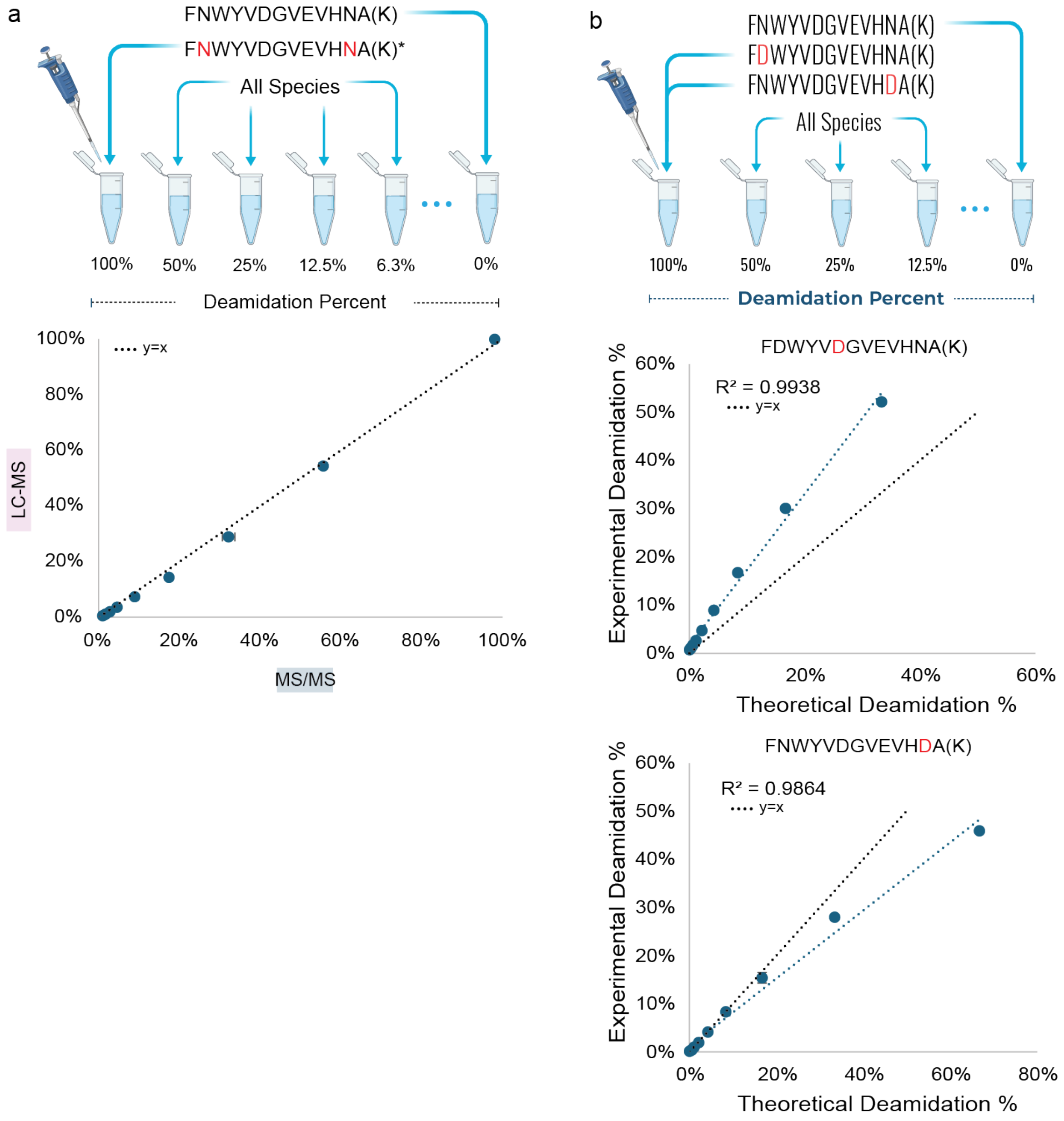


**a**, Titration of $^{13}C$ labeled native and deamidated FNWYVDGVEVHNA(K) peptides to make solution from 100% deamidated to 0% deamidated in a background of NIST mAb peptides with comparisons of LC-MS and RaPiD-mAb-MS analysis of the spike-in mixtures showing high correlation between the methods (n=3). **b**, Site specific titration of each deamidation site on $^{13}C$ labeled FNWYVDGVEVHNA(K) peptide.

**Extended Data Figure 4: AbbVie antibody-drug conjugate (AB095-PZ) analysis by MS1 and DDA MS/MS: THTCPPCPAPELLGGPSVFLFPPKPK peptide.**

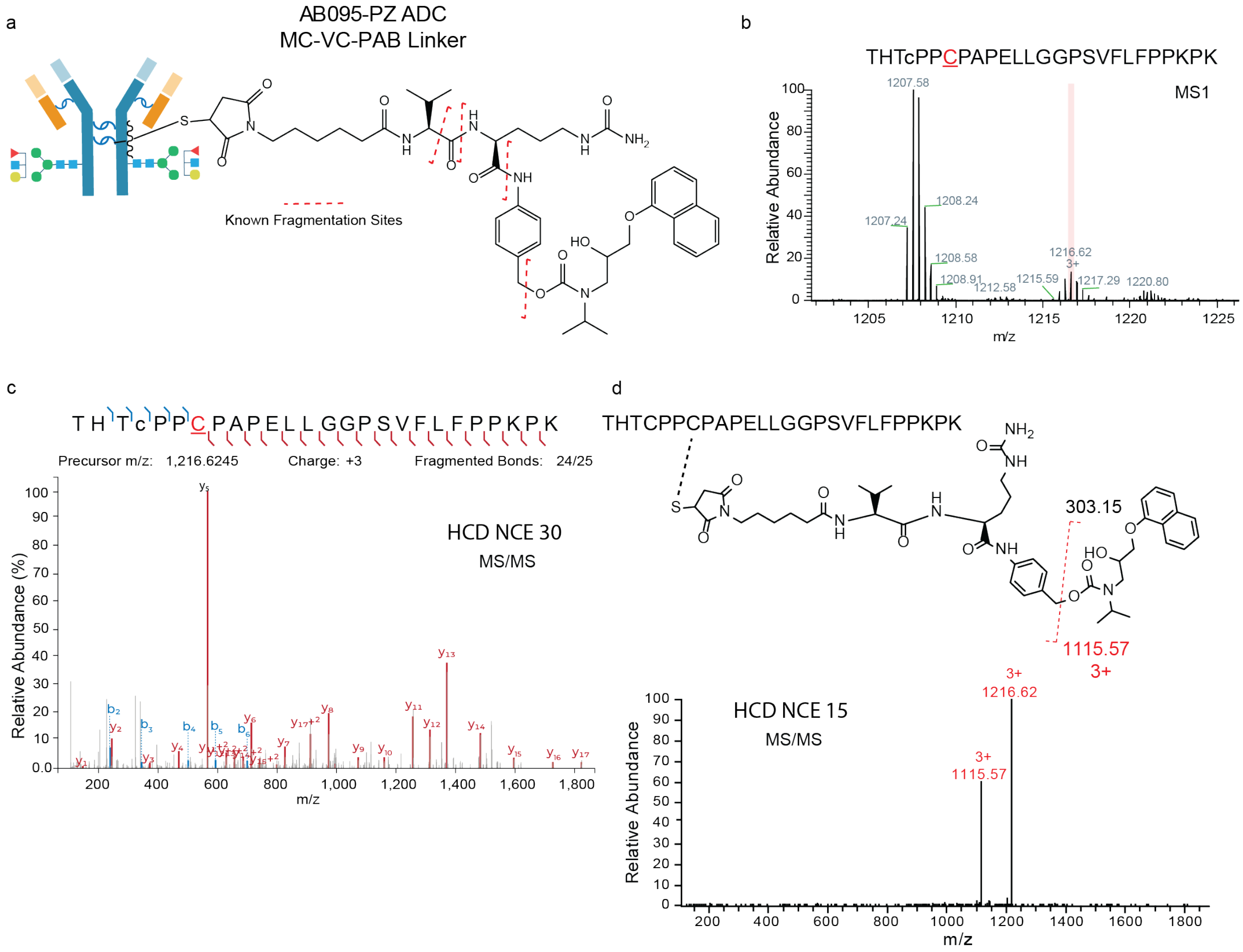


**a**, Linker and drug structure for AB095-PZ ADC. **b,** MS1 region of RaPiD-mAb-MS infusion data highlighting the isotope distribution of an intact peptide with MC-VC-PAB linker and pseudo drug, as confirmed by accurate mass. **c**, MS/MS spectrum of the identified MS1 peak at m/z value 1216.62 (THTCPPCPAPELLGGPSVFLFPPKPK + linker and drug; (Cys233)). This MS/MS spectrum both confirms the accurate MS1 identification and allows for localization of the drug and linker to the underlined cysteine. **d**, Lower energy HCD scan demonstrates known fragmentation of the linker releasing the pseudo drug as a neutral loss.

**Extended Data Figure 5: AbbVie antibody-drug conjugate (AB095-PZ) analysis by MS1 and DDA MS2: Small Peptides.**

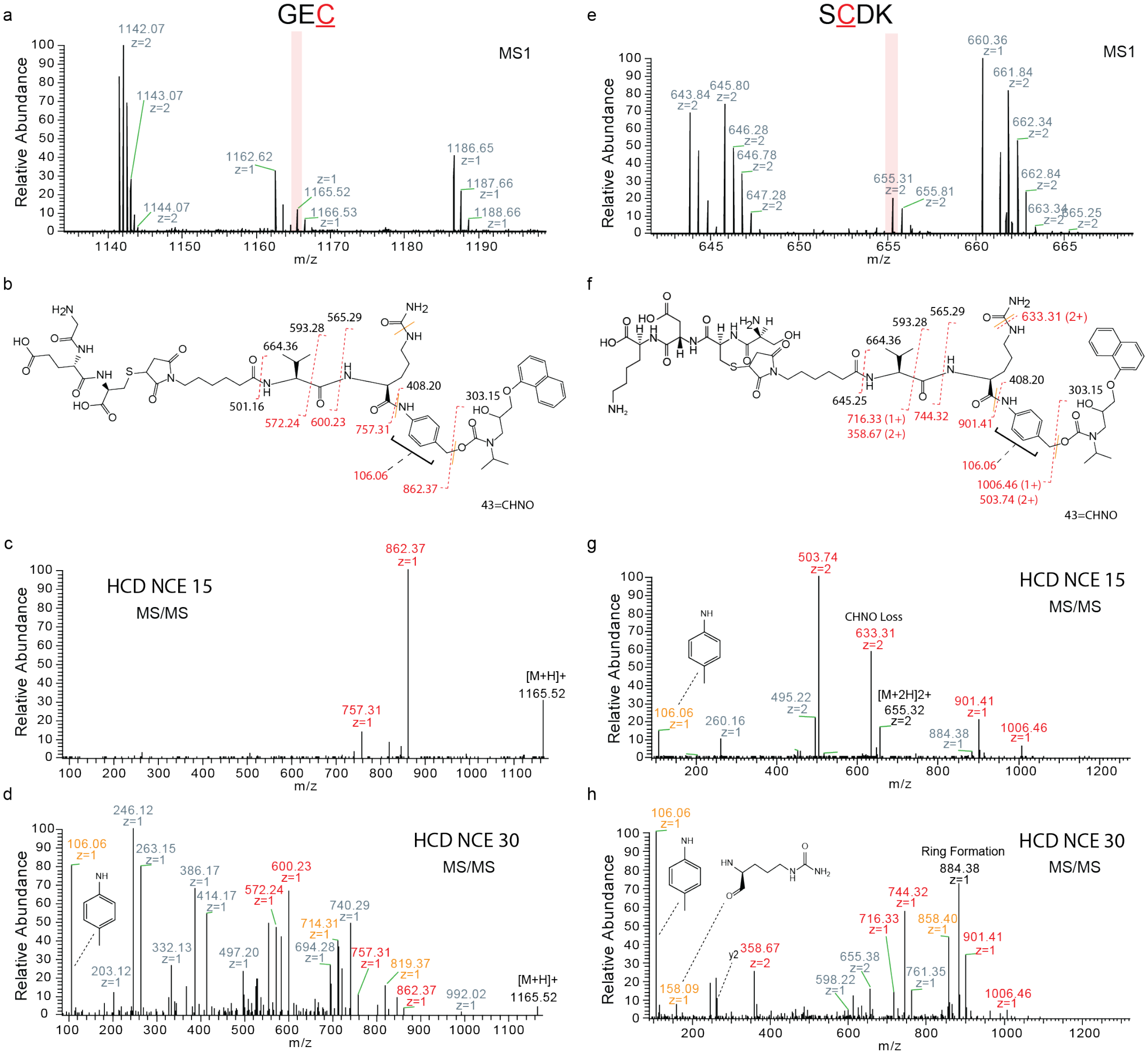


**a,** MS1 region of RaPiD-mAb-MS infusion data highlighting the isotope distribution of an intact GEC peptide with MC-VCPABC linker and pseudo drug, as confirmed by accurate mass. **b**, Peptide, Linker and drug structure for AB095-PZ ADC. **c**, Low energy MS/MS spectrum of the identified MS1 peak at m/z value 1162.52 (GEC + linker and drug) demonstrating known fragmentation. **d**, High energy MS/MS spectrum of the identified MS1 peak at m/z value 1162.52 (GEC + linker and drug). This MS/MS spectrum both confirms the accurate MS1 identification and allows for localization of the drug and linker to the underlined cysteine. **e-h**, Shows the same analysis but for SCDK peptide with linker and drug. Red dotted lines denote fragments consistent with a single fragmentation event, whereas orange solid lines denote fragments that require two fragmentation events.

**Extended Data Figure 6: Global RaPiD-mAb-MS and LC-MS peptide mapping comparison for NIST mAb.**

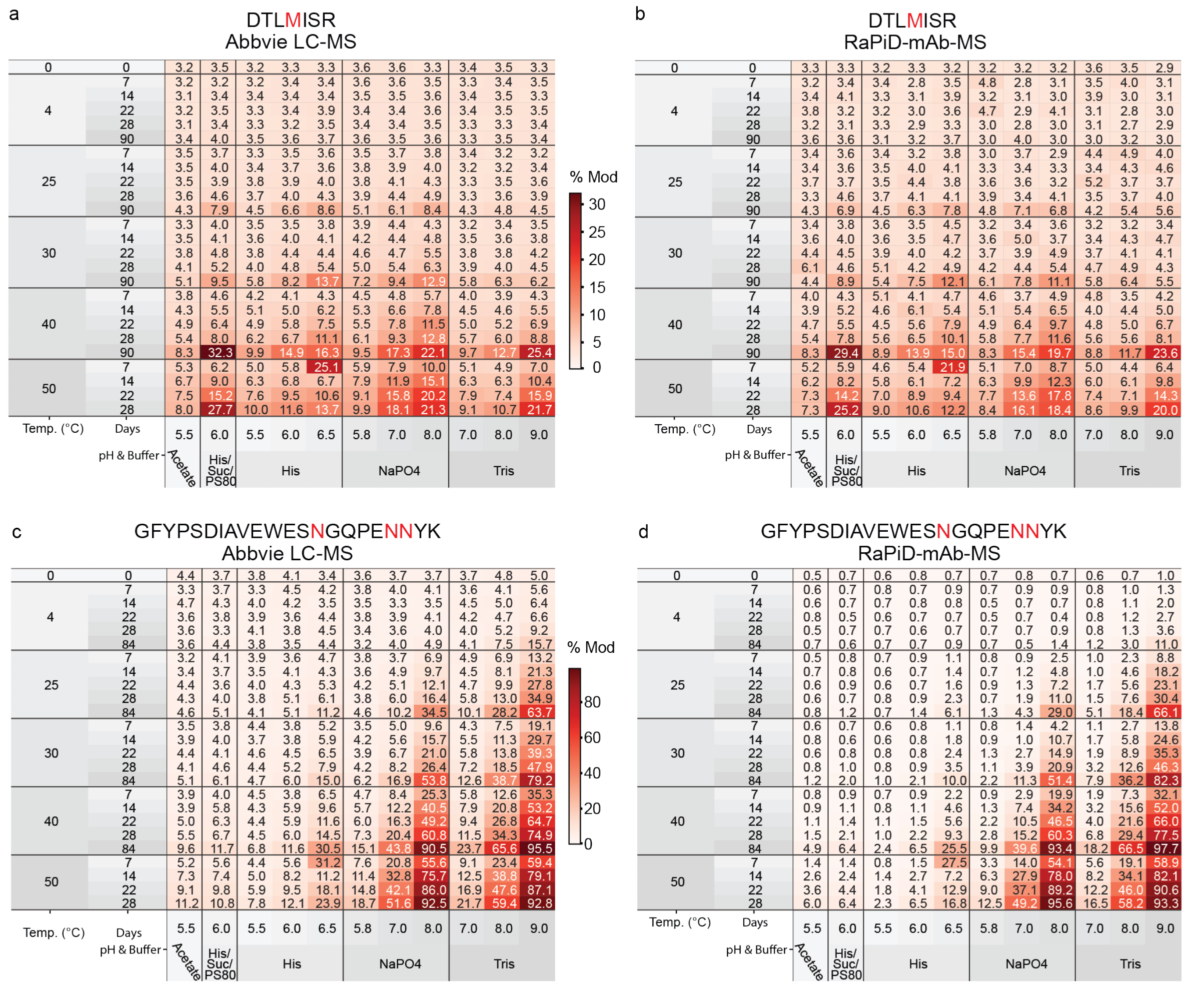


Heatmaps showing the relative modification of the DTMLISR peptide across 275 samples from (**a**) AbbVie LC-MS and (**b**) RaPiD-mAb-MS data, and of the deamidated PENNYK peptide from (**c**) AbbVie LC-MS and (**d**) RaPiD-mAb-MS data.

**Extended Data Figure 7: RaPiD-mAb-MS and LC-MS peptide mapping comparison of other modification sites.**

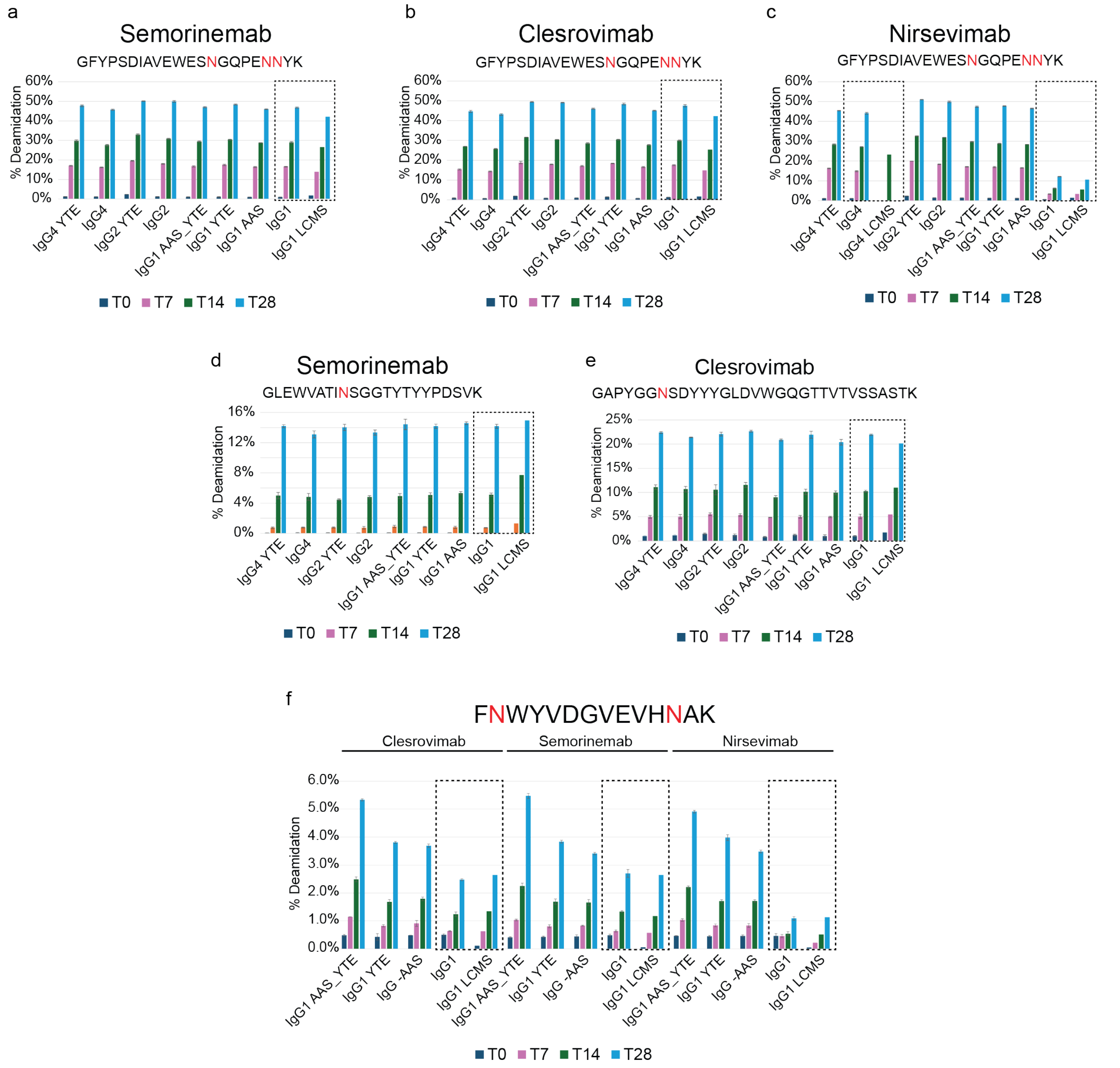


**a-f**, Comparison of a variety of mAbs across time points for deamidation with dashed box showing samples analyzed by both RaPiD-mAb-MS and LC-MS. All values shown represent peptide-level (summed) occupancies rather than residue-localized measurements.

#### Extended Data Figure 8: Heatmap for oxidation conditions.

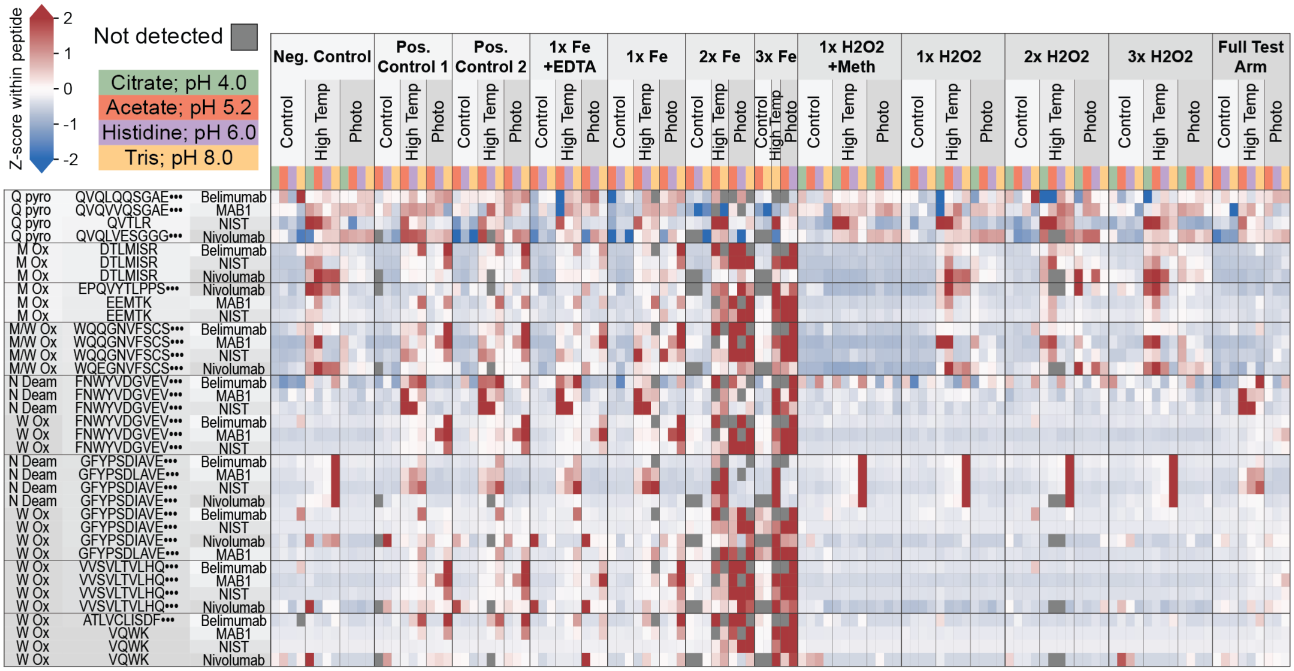


Heatmap showing the RaPiD-mAb-MS measured quantitative differences for the conserved modifications sites across four antibodies for the 144 oxidation experiment conditions.

**Extended Data Figure 9: Bar plots of NIST mAb peptides from oxidation conditions with single methionine and tryptophan present as well as the combination in one peptide.**

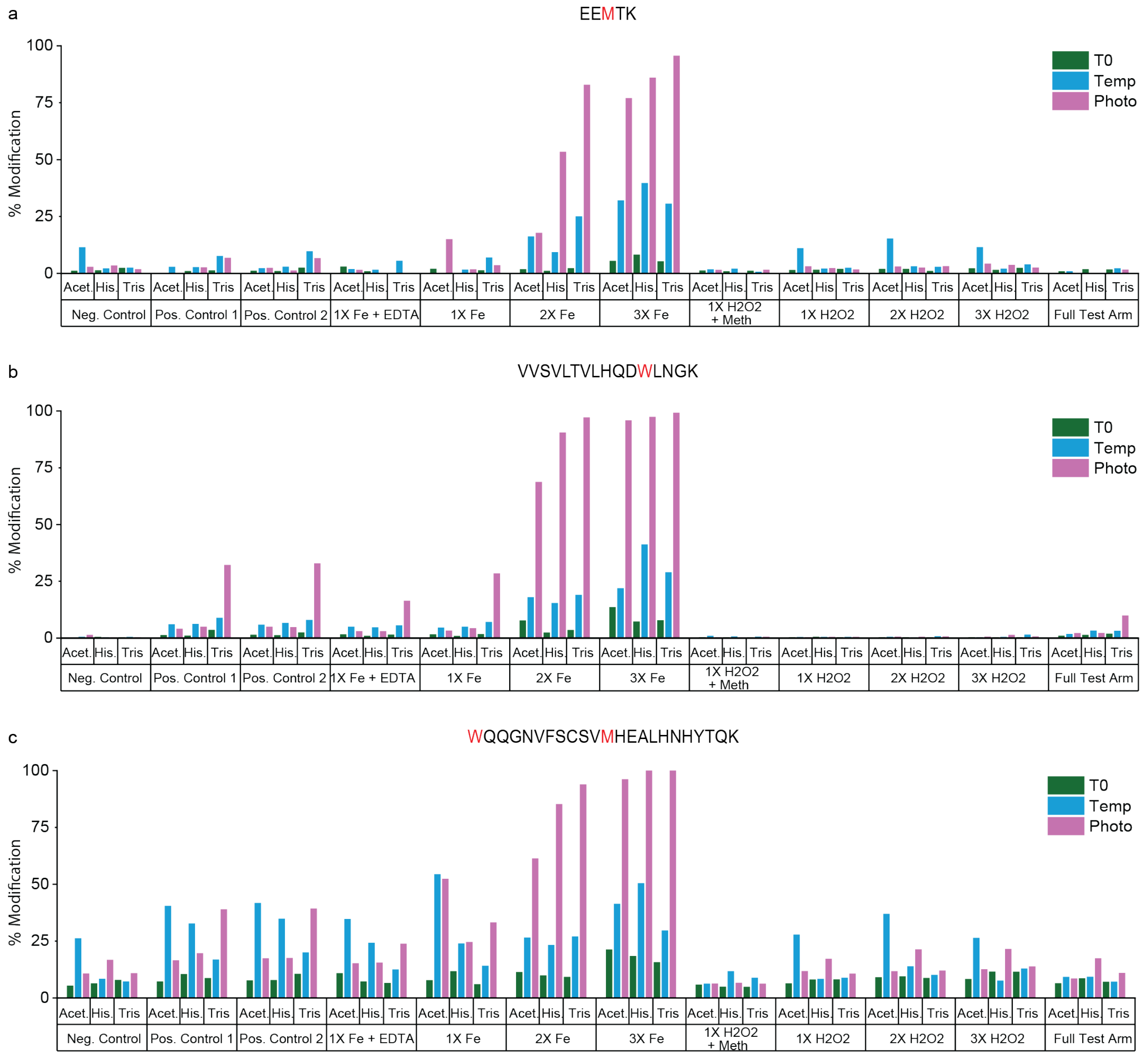


Bar plots across 3 different peptides from NIST mAb including (**a**) EEMTK, (**b**) VVSVLTVLHQDWLNGK, and (**c**) WQQGNVFSCSVMHEALHNHYTQK from the oxidation conditions. All values shown represent peptide-level (summed) occupancies rather than residue-localized measurements.

**Extended Data Figure 10: Heatmap for NIST mAb for both buffer and oxidation conditions and effect of tested stressors, excipients, buffer and pH on average modification percent on 16 modification sites in NIST mAb.**

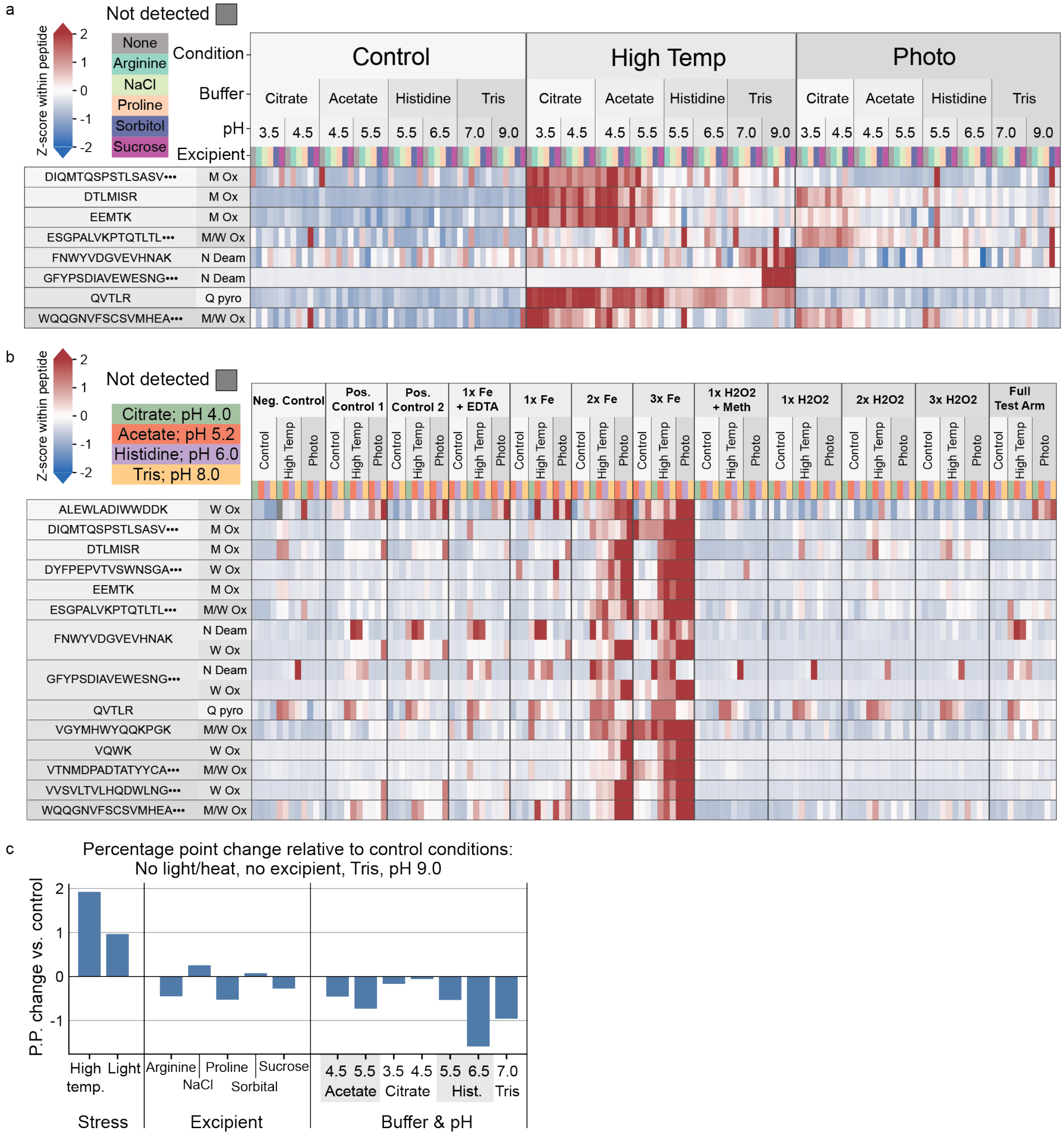


**a**, Heat map of NIST mAb peptides with modifications present for buffer experiments. **b**, Heat map of NIST mAb peptides with modifications present for oxidation experiments. **c**, Percentage point changes given here are effect sizes from a linear model trained on peptide data across all conditions across modification sites. A positive bar means that the additive causes modifications to increase.

**Extended Data Figure 11: Accurate isoAsp quantitation in stressed NIST mAb by DI-MS and LC-MS, benchmarked across a native-to-aged dilution series.**

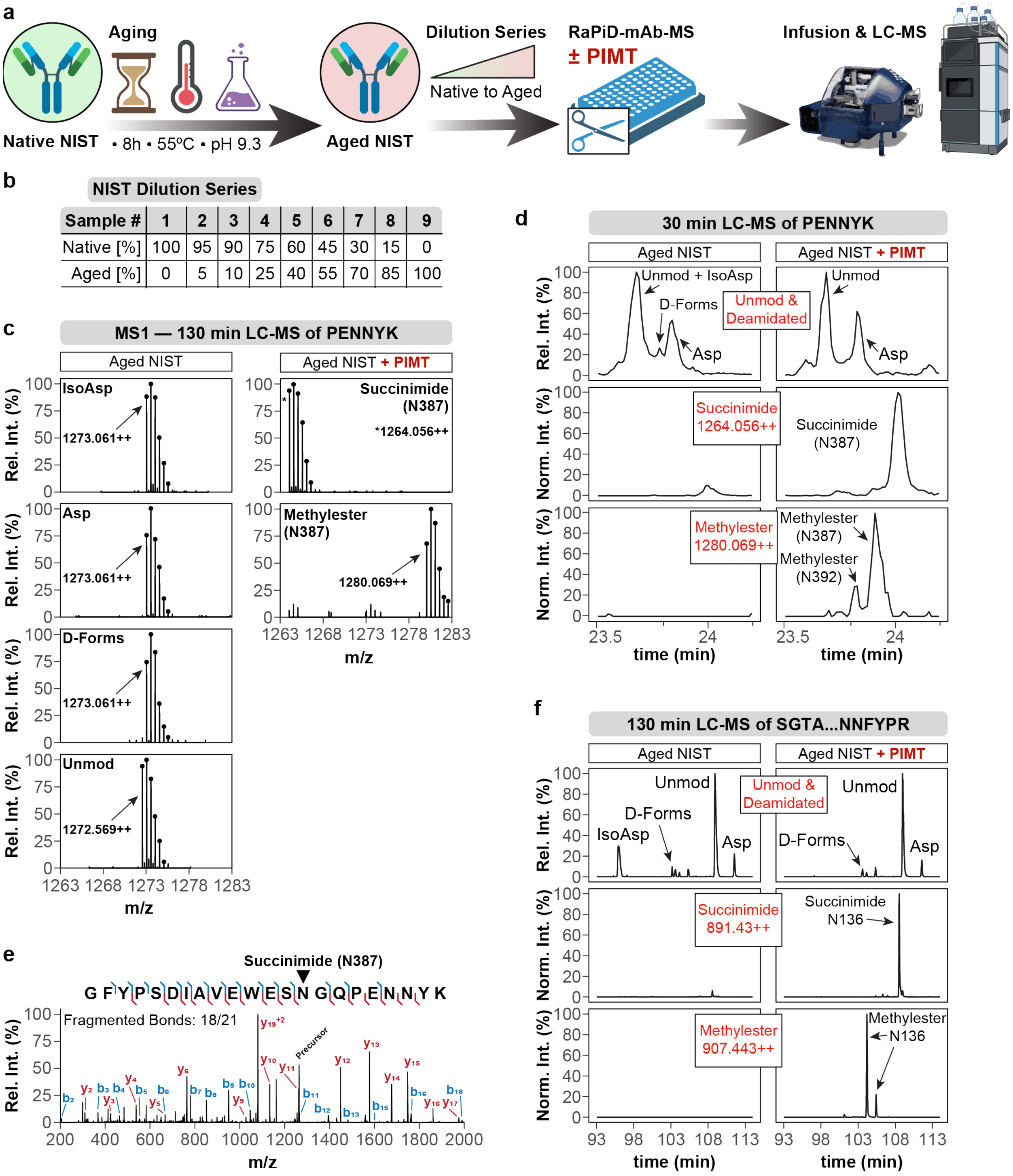


| Sample # | 1 | 2 | 3 | 4 | 5 | 6 | 7 | 8 | 9 |
|---|---|---|---|---|---|---|---|---|---|
| Native [%] | 100 | 95 | 90 | 75 | 60 | 45 | 30 | 15 | 0 |
| Aged [%] | 0 | 5 | 10 | 25 | 40 | 55 | 70 | 85 | 100 |

**a,** NIST mAb aging workflow and downstream analysis. Native NIST mAb was subjected to combined thermal and high-pH stress (55 °C, pH 9.3, 8 h) to generate an aged NIST mAb sample, followed by preparation of a native-to-aged dilution series. Samples were then processed within the RaPiD-mAb-MS workflow with or without PIMT treatment and analyzed by direct infusion MS and LC-MS. **b,** NIST mAb dilution series. Native and Aged NIST mAb were mixed at defined ratios to generate a nine-point dilution series (Samples 1-9), spanning 0-100% Aged NIST mAb (and corresponding 100-0% Native NIST mAb) as indicated. **c,** MS1 spectra at peak apex for chromatographic features of PENNYK deamidation-related species in aged NIST mAb (+-PIMT). Representative MS1 spectra (m/z 1263-1283) were extracted at the apex of each LC peak shown in Fig. 7c for the PENNYK peptide during a 130 min LC-MS run. Spectra are shown for aged NIST mAb without PIMT (left: IsoAsp, Asp, D-forms, and unmodified windows) and aged NIST mAb with PIMT (right: succinimide and methylester windows at N387). The MS1 spectra confirm the expected mass equivalence of the isobaric deamidation-related forms (IsoAsp/Asp/D-forms) and thus support the overall peak assignments, but do not by themselves distinguish among these isobars. The identities of the individual LC peaks were established based on peptide elution behavior in supporting experiments. **d,** short-gradient LC-MS traces for PENNYK in aged NIST mAb (+-PIMT). Extracted-ion chromatograms (XICs) for PENNYK acquired using a short LC-MS gradient (30 min) are shown for aged NIST mAb without PIMT (left) and with PIMT treatment (right), using the same trace layout as in Fig. 7c. Under the short-gradient conditions, the unmodified and IsoAsp-associated signals are not baseline-resolved and appear as a merged feature in the -PIMT sample. In the +PIMT sample, the succinimide and methylester products remain detectable in their respective windows due to their mass shifts. **e,** Representative MS2 spectrum for succinimide forms of PENNYK acquired using a targeted PRM scan (related to Figure 7d). **f,** long-gradient LC-MS traces for SGTASVVCLLNNFYPR in aged NIST mAb (+-PIMT). Related to panel d and to Fig. 7c.

### Supplementary Information

#### Supplementary Figures

Supplementary Figure. 1: Segmented MS1 scan of NIST mAb.

Supplementary Figure. 2: MS performance with changing concentration, MS1 AGC target, and MS1 resolution.

#### Supplementary Tables

Supplementary Table 1: Buffer conditions for the J&J samples provided include concentration, buffer, pH, and excipient.

Supplementary Table 2: Oxidation conditions for the J&J samples including concentration, buffer, pH, and excipient (H2O2, Fe, Meth, EDTA).

Supplementary Table 3:
Additional information on each excipient added for oxidation conditions.

Supplementary Table 4: MS instruments used for direct infusion MS and/or LC-MS datasets across figures and panels.

Supplementary Table 5: Linearity ($R^2$) of RaPiD-mAb-MS quantitation in the 0-10% occupancy range for panels in Figures 2 and 3.

Supplementary Table 6: Detected NIST mAb peptides and MS1 features supporting sequence coverage.

#### Supplementary Figure 1: Segmented MS1 scan of NIST mAb.

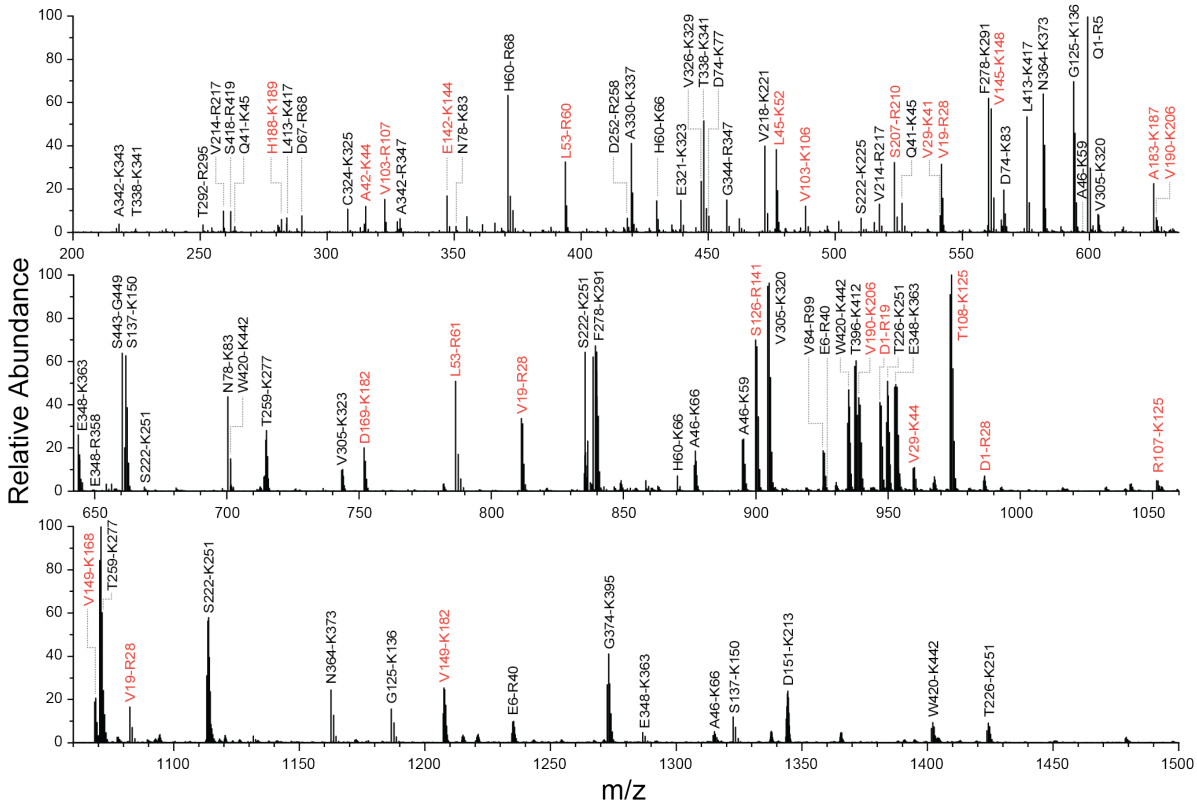


Segmented MS1 scan of NIST mAb with labels for the heavy (black) and light (red) chain. This sample has an overall sequence coverage of 96% by hand with a % total ion chromatogram (TIC) explained of 96%. This was determined by calculating the relative percent of each isotope cluster to the base peak with a threshold cut-off of 0.5%. The full list of detected peptides (including observed charge states and m/z values) is provided in **Supplementary Table 6**.

**Supplementary Figure 2: MS performance with changing concentration, AGC target, and MS1 resolution.**

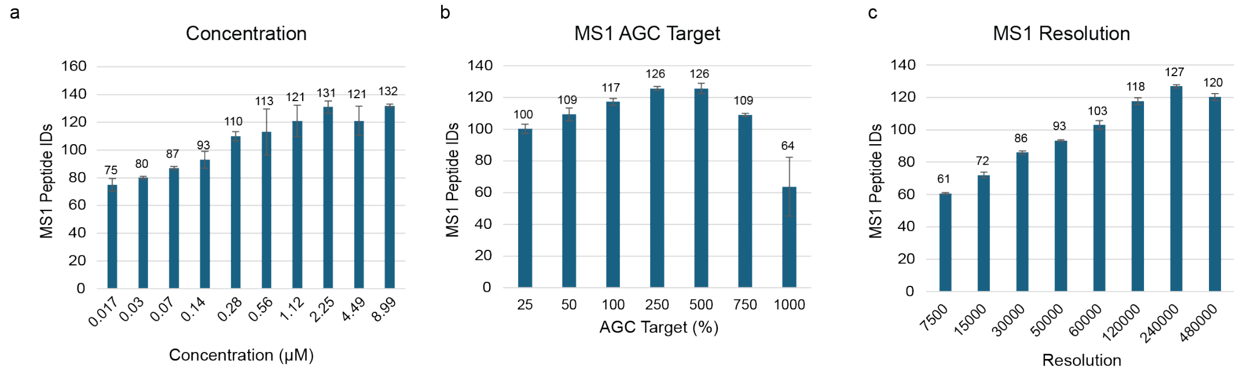


**a**, Effects of concentration on number of unique peptides identified with a S/N cut-off of 3 (n=3). **b**, Ideal AGC target determined for 480K MS1 resolving power with both 250% and 500% showing similar results (n=3). **c**, MS1 resolution showing similar results from 120K-480K resolving power (n=3).

**Supplementary Table 1: Buffer conditions for the J&J samples provided include concentration, buffer, pH, and excipient.**

| Buffer | | | |
|---|---|---|---|
| **mg/mL** | **Buffer** | **pH** | **Excipient** |
| 2 | Citrate | 3.5 | None |
| 2 | Citrate | 3.5 | Proline |
| 2 | Citrate | 3.5 | Arginine |
| 2 | Citrate | 3.5 | NaCl |
| 2 | Citrate | 3.5 | Sucrose |
| 2 | Citrate | 3.5 | Sorbitol |
| 2 | Citrate | 4.5 | None |
| 2 | Citrate | 4.5 | Proline |
| 2 | Citrate | 4.5 | Arginine |
| 2 | Citrate | 4.5 | NaCl |
| 2 | Citrate | 4.5 | Sucrose |
| 2 | Citrate | 4.5 | Sorbitol |
| 2 | Histidine | 5.5 | None |
| 2 | Histidine | 5.5 | Proline |
| 2 | Histidine | 5.5 | Arginine |
| 2 | Histidine | 5.5 | NaCl |
| 2 | Histidine | 5.5 | Sucrose |
| 2 | Histidine | 5.5 | Sorbitol |
| 2 | Histidine | 6.5 | None |
| 2 | Histidine | 6.5 | Proline |
| 2 | Histidine | 6.5 | Arginine |
| 2 | Histidine | 6.5 | NaCl |
| 2 | Histidine | 6.5 | Sucrose |
| 2 | Histidine | 6.5 | Sorbitol |

| Buffer | | | |
|---|---|---|---|
| **mg/mL** | **Buffer** | **pH** | **Excipient** |
| 2 | Acetate | 6.5 | None |
| 2 | Acetate | 4.5 | Proline |
| 2 | Acetate | 4.5 | Arginine |
| 2 | Acetate | 4.5 | NaCl |
| 2 | Acetate | 4.5 | Sucrose |
| 2 | Acetate | 4.5 | Sorbitol |
| 2 | Acetate | 5.5 | None |
| 2 | Acetate | 5.5 | Proline |
| 2 | Acetate | 5.5 | Arginine |
| 2 | Acetate | 5.5 | NaCl |
| 2 | Acetate | 5.5 | Sucrose |
| 2 | Acetate | 5.5 | Sorbitol |
| 2 | Tris | 7 | None |
| 2 | Tris | 7 | Proline |
| 2 | Tris | 7 | Arginine |
| 2 | Tris | 7 | NaCl |
| 2 | Tris | 7 | Sucrose |
| 2 | Tris | 7 | Sorbitol |
| 2 | Tris | 9 | None |
| 2 | Tris | 9 | Proline |
| 2 | Tris | 9 | Arginine |
| 2 | Tris | 9 | NaCl |
| 2 | Tris | 9 | Sucrose |
| 2 | Tris | 9 | Sorbitol |

Buffer Conditions: 3 Plates x 48 Conditions=144 Unique Combinations. 3 Plates=(control or no temp, Photo=30Whrs, Temp=40$^{o}$C for 14 days).

**Supplementary Table 2: Oxidation conditions for the J&J samples including concentration, excipients ($H_2O_2$, Fe, Meth, EDTA).**

| Oxidation | | | | | |
|---|---|---|---|---|---|
| mg/mL | H2O2 | Fe | Meth | EDTA | Note |
| 2 | – | – | – | – | Negative Control |
| 2 | + | – | – | – | H2O2 affects |
| 2 | – | + | – | – | Fe Affects |
| 2 | + | + | – | – | Positive control -1 |
| 2 | + | – | + | – | H2O2/Meth treatment |
| 2 | – | + | – | + | Fe/EDTA Treatment |
| 2 | + | + | + | + | Full test arm |
| 2 | 2x | – | – | – | 2x H2O2 affects |
| 2 | 3x | – | – | – | 3x H2O2 affects |
| 2 | – | 2x | – | – | 2x Fe Affects |
| 2 | – | 3x | – | – | 3x Fe Affects |
| 2 | + | + | – | – | Positive control -2 |

Buffers for the above conditions include Citrate (pH 4.0), Acetate (pH 5.2), Histidine (pH 6.0), and Tris (pH 8.0) making for 48 unique conditions.

**Supplementary Table 3: Additional information on each excipient added for oxidation conditions.**

| Additional Information | |
|---|---|
| – | not added |
| + | added |
| 2x/3x | 2x/3x indicate 2- or 3-fold higher amounts relative to the standard "+" condition. |
| [redacted] | [redacted] |
| [Meth], mg/mL | 1 |
| [EDTA], ug/mL | 20 |
| [H2O2], ppb | 50 |
| [Fe], ppb | 50 |

Oxidation Conditions: 3 Plates x 48 Conditions=144 Unique Combinations. 3 Plates = (Control or no temp, Photo=30Whrs, Temp=40°C for 14 days).

**Supplementary Table 4. MS instruments used for direct infusion MS and/or LC-MS datasets.**

| Figure | DI-MS | LC-MS |
|---|---|---|
| Figure 1 | Eclipse | — |
| Figure 2a-c | Eclipse | Eclipse |
| Figure 2d | Astral | — |
| Figure 3 | Astral | Ascend |
| Figure 4 | Eclipse | — |
| Figure 5a-f | Astral | Astral |
| Figure 5g | Eclipse | Eclipse |
| Figure 6 | Astral | — |
| Figure 7 | Ascend | Ascend, Exploris 240, Astral |
| Extended Data Figure 1 | Astral | — |
| Extended Data Figure 2 | Eclipse | Eclipse |
| Extended Data Figure 3 | Astral | Ascend |
| Extended Data Figure 4 | Eclipse | — |
| Extended Data Figure 5 | Eclipse | — |
| Extended Data Figure 6 | Astral | Q-Exactive Plus (AbbVie) |
| Extended Data Figure 7 | Eclipse | Eclipse |
| Extended Data Figure 8 | Astral | — |
| Extended Data Figure 9 | Astral | — |
| Extended Data Figure 10 | Astral | — |
| Extended Data Figure 11 | — | Exploris 240, Astral |
| Supplementary Figure 1 | Eclipse | — |
| Supplementary Figure 2 | Eclipse | — |

**Supplementary Table 5. Linearity ($R^2$) of RaPiD-mAb-MS quantitation in the 0-10% occupancy range for panels in Figures 2 and 3.**

| Figure panel | Peptide | $R^2$ (0–10%) |
|---|---|---|
| 2b | DTLMISR | 0.999 |
| 2b | WQQGNVFSCSVMHEALHNHYTQK | 0.999 |
| 2b | FNWYVDGVEVHNAK | 0.999 |
| 2b | VVSVLTVLHQDWLNGK | 0.998 |
| 3d | GFYPSDIAVEWESNGQPENNYK | 0.855 |
| 3e | GFYPSDIAVEWESDGQPENNYK | 0.983 |
| 3e | GFYPSDIAVEWESNGQPEDNYK | 0.997 |
| 3e | GFYPSDIAVEWESNGQPENDYK | 0.974 |

Note: In the peptide sequences listed, the modified (relevant) amino acid are shown in red.
